# Reassembly of a tropical rainforest ecosystem: A new chronosequence in the Ecuadorian Chocó tested with the recovery of tree attributes

**DOI:** 10.1101/2024.03.21.586145

**Authors:** Sebastián Escobar, Felicity L. Newell, María-José Endara, Juan E. Guevara-Andino, Anna R. Landim, Eike Lena Neuschulz, Ronja Nußer, Jörg Müller, Karen M. Pedersen, Matthias Schleuning, Constance J. Tremlett, Edith Villa-Galaviz, H. Martin Schaefer, David A Donoso, Nico Blüthgen

## Abstract

From hunting and foraging to clearing land for agriculture, humans modify forest biodiversity, landscapes, and climate. Forests constantly undergo disturbance–recovery dynamics and understanding them is a major objective of ecologists and conservationists. Chronosequences are a useful tool for understanding global restoration efforts. They represent a space-for-time substitution approach suited for the quantification of the *resistance* of ecosystem properties to withstand disturbance and the *resilience* of these properties until reaching pre-disturbance levels. Here we introduce a newly established chronosequence with 62 plots (50 ⍰ 50 m) in active cacao plantations and pastures, early and late regeneration, and mature old-growth forests, across a 200 km2 area in the extremely wet Chocó rainforest. Our chronosequence covers by far the largest total area of plots compared to others in the Neotropics. Plots ranged from 159–615 masl in a forested landscape with 74 ± 2.8 % forest cover within a 1-km radius including substantial old-growth forest cover. Land-use legacy and regeneration time were not confounded by elevation. We tested how six forest structure variables (maximum tree height and DBH, basal area, number of stems, vertical vegetation heterogeneity, and light availability), aboveground biomass (AGB), and rarefied tree species richness change along our chronosequence. Forest structure variables, AGB, and tree species richness increased with regeneration time and are predicted to reach similar levels to those in old-growth forests after ca. 30–116, 202, and 108 yrs, respectively. Compared to previous work in the Neotropics, old-growth forests in Canandé accumulate high AGB that takes one of the largest time spans reported until total recovery. Our chronosequence comprises one of the largest tree species pools, covers the largest total area of regenerating and old-growth forests, and has higher forest cover than other Neotropical chronosequences. Hence, our chronosequence can be used to determine the time for recovery and stability (resistance and resilience) of different taxa and ecosystem functions, including species interaction networks. This integrative effort will ultimately help to understand how one of the most diverse forests on the planet recovers from large-scale disturbances.

## 1. Introduction

Tropical forests are highly threatened by deforestation, degradation, and climate change. Every year, some 4.5 million ha of old-growth forests are logged and mostly replaced by pastures and crops (FAO, 2020). This disturbance entails a reduction in species diversity and in the complex interaction networks knitted between them (Poorter et al., 2021a, b). The ongoing ‘UN decade of ecosystem restoration’ underscores a global urgency to mitigate losses in ecosystems and bend the curve of biodiversity decline, but requires a sound understanding of ecological dynamics (Cooke et al., 2019). Knowledge of the active restoration and natural recovery potential of tropical forests is not only needed for ecological studies, but also for conservation and restoration implementers. Studying chronosequences covering different stages of natural forest regeneration allows for a deeper understanding of succession and the extent to which secondary forests recover (Figure 1; Chazdon et al., 2007). Here, we introduce the theoretical background and spatial design of a well-resolved chronosequence in a Neotropical rainforest that allows us to study the recovery of plant and animal communities and the interactions that maintain this highly diverse ecosystem. We analyse baseline data as a case study to show how six measures of forest structure, aboveground biomass (AGB), and tree species richness change with regeneration time along with other abiotic variables.

**Figure 1.**
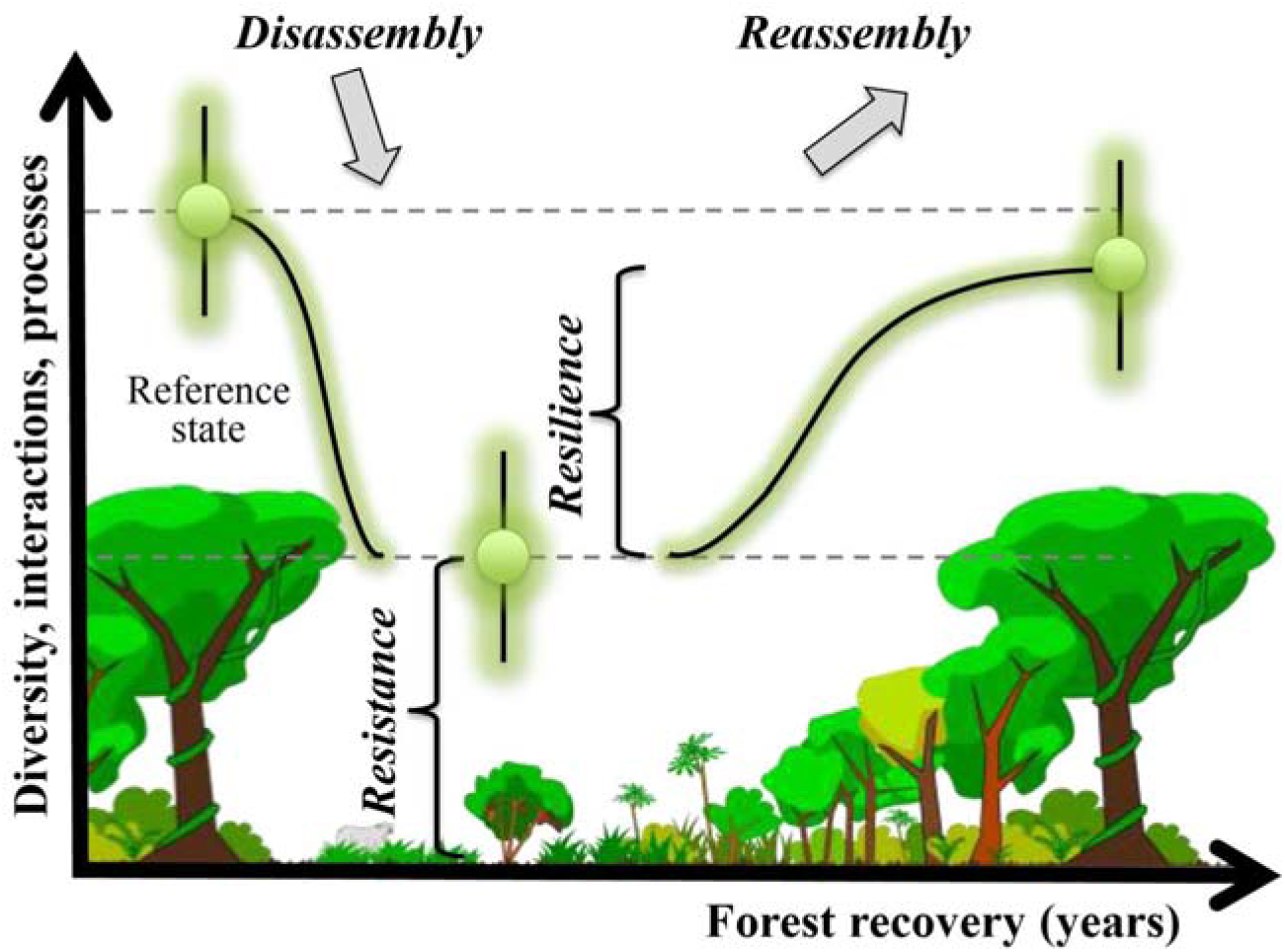
Conceptual framework of ecosystem dis- and reassembly, showing the proposed dynamics of diversity, interactions, and related ecosystem processes. The first step describes the reduction or disassembly of old-growth forests’ properties, following disturbance by deforestation and subsequent land use as pasture or plantation. The amount of diversity, interactions, and processes that withstands disturbance and remains present during agricultural use defines the resistance of that property. The recovery of forests or reassembly of communities and networks starts as soon as the utilization as pasture or plantation stops. We generally predict a continuous increase of diversity, network complexity, and process rates with forest recovery until some saturation level (if there is no support for a hump-shaped trajectory for alpha diversity as predicted by the intermediate disturbance hypothesis). The level reached after a given time of recovery may be lower (incomplete reassembly) or similar to the average level of old-growth forests. The speed and extent of recovery represent the resilience of the forest ecosystem.

Tropical forests are key biomes because they harbour over half of the global biodiversity (Pimm & Raven, 2000) and account for one-third of primary productivity on the land surface by sequestering 1.7 Gt of carbon per year (Malhi, 2012; Harris et al., 2021). Tropical ecosystems are inherently dynamic, going through stages of natural and anthropogenic **disturbances** (see **Glossary** for definitions of key terms). Recovery occurs at various spatial and temporal scales and provides services and renewable resources for more than one billion people (Ghazoul et al., 2015; Lewis et al., 2015; Ghazoul & Chazdon, 2017). Despite their importance, half of tropical forests had already disappeared by the beginning of the twentieth century (Wright, 2005), and deforestation continues at high levels today (FAO, 2020).

Worldwide demand for palm oil, soybeans, rubber, timber, and other cash crop products contributed to unprecedented levels of conversion of tropical forests during the last decades (Watson et al., 2018). Shifting agriculture is one of the primary drivers of forest loss in tropical regions (Curtis et al., 2018). The disturbance of tropical forests for agricultural purposes can promote edge effects and the isolation of forest patches (Malhi et al. 2014; Wanyama et al. 2023). Distance to source populations and the amount of forest left at different radii are critical factors influencing how animals mediate key ecosystem functions such as seed dispersal, pollination, host plant defense, predation, parasitism, and decomposition (Lundberg & Moberg, 2003; Tylianakis & Morris, 2017). Therefore, forest conversion can lead to profound changes in forest structure and composition, affecting different dimensions of biodiversity and their functional consequences. These changes may include a strong decrease in biomass and species diversity losses in plant and animal communities (Longo et al., 2016; Baccini et al., 2017; Pfeifer et al., 2017; Matricardi et al., 2020). For instance, only 59 % of plant or animal species occur in disturbed and secondary habitats altogether when compared to old-growth forests (Alroy, 2017). Thus, for the conservation and long-term permanence of tropical forests, it is fundamental to understand how they recover after disturbance, which factors promote recovery, and at which speed they operate (Chazdon et al., 2009; Gardner et al., 2009).

### 1.1 Resistance, resilience, and functional recovery of tropical forests

Measuring forest recovery is challenging, and it often requires a definition of stability in time. Stability can be defined for either single properties of a system (e.g., population density, diversity, network metric, or a mean process rate) or for multiple dimensions, including variation within and among properties (Pimm, 1984; Donohue et al., 2013; Hillebrand et al., 2018). Several concepts of **stability** exist, including resistance and resilience (Van Meerbeek et al., 2021). **Resistance** (Figure 1) mirrors the proportion of species, communities, interactions, and processes that are sustained following **perturbation** and thus persist in an altered ecosystem – facilitated by properties to resist immediate impact (pulse perturbation) and/or changes in environmental conditions in continued disturbance regimes (press perturbation; Pimm, 1984; McCann, 2000; White et al., 2020). **Resilience** can be defined as the recovery rate of the amount lost to a stable level (often close to an ‘equilibrium’) found in an undisturbed reference ecosystem, i.e., the speed of **reassembly** or its completeness after a certain time span (Holling et al., 1973; Folke et al., 2004; Pimm et al., 2019). **Recovery time** describes the span from the time disturbance halts until recovery is reached; and can be accelerated both by high resistance and by high resilience independently (Pimm et al., 2019).

Species communities and their ecosystem functions are known to vary in their resistance and resilience. Some taxa such as trees almost completely vanish in agricultural landscapes while others, such as ants, become more abundant (Dunn et al., 2004; Rozendaal et al., 2019; Hoenle et al., 2022). Measurements of recovery time across different systems and taxa, including changes in trait rules and species interactions in time, thus contribute to our understanding of variation in resilience. However, these variables have seldom been measured *in situ*. A recent global meta-analysis (Biggs et al., 2020) found only 15 studies (none in tropical forests) that tested whether functional redundancy of traits improves stability in terrestrial ecosystems. Therefore, the resistance and resilience of ecological communities to ecosystem conversion are still poorly assessed, hampering our understanding of their potential recovery.

### 1.2 Chronosequences substitute time with space

Most contemporary assessments of tropical forest resilience, in terms of structure and species diversity, are based on **chronosequences**. Chronosequences allow the study of forest succession (Chazdon et al., 2007) while examining forest resilience and projecting its trajectory. Generally, the trajectory of the resilience of diversity (or network complexity) may either be linear (or saturating) or non-linear (hump-shaped, unimodal, etc.). The ‘**intermediate disturbance hypothesis**’ (Grime, 1973; Connell, 1978) predicts a hump-shaped trajectory, with a diversity peak at intermediate stages of recovery after disturbance and/or at an intermediate disturbance level or frequency. Surprisingly, only a few tests of this influential hypothesis have proved its validity for tree diversity, for which it was proposed (Hubbell et al., 1999; Molino & Sabatier, 2001). In addition, non-linearity has not been explicitly considered and tested in published chronosequences of reassembling animal communities (e.g., Dunn, 2004; Crouzeilles et al., 2016 and references therein). The recovery of forests is generally expected – but rarely shown – to be fastest for species diversity, intermediate for species composition, and slowest for complex species interaction networks (Acevedo-Charry & Aide, 2019; Poorter et al., 2021a).

### 1.3 The Chocó rainforest

The Chocó rainforest is a biodiversity hotspot that is highly threatened by deforestation. The Chocó is located in northwestern South America to the west of the Ecuadorian and Colombian Andes and in Panamá, occupying only 0.2% of Earth’s land surface (Myers et al., 2000; Pérez-Escobar et al., 2019). However, it is considered the world’s ninth most biodiverse hotspot, harboring ∼11,000 plant species (3% of the total), 25% of which are endemic (Gentry, 1982; Christenhusz et al., 2017). The Chocó is a highly threatened habitat that has been destroyed at very high rates. The particularly high deforestation rate in this area has been quantified by satellite images available since 1986 (Gonzalez-Jaramillo et al., 2016), and until 2015, only a small amount of the old-growth forest (<11%) was left (Fagua et al. 2019).

## 2. The Reassembly Project

Given the extensive deforestation and natural regeneration in the Chocó, this ecosystem is ideal to understand resistance, resilience, and recovery of a tropical rainforest. Here we present the design of a collaborative German-Ecuadorian Research Unit “*Reassembly of species interaction networks”* aiming to unravel to what extent, and how fast a rainforest can re-establish itself after deforestation. This includes the diverse plant and animal communities, their complex interaction networks, and relevant ecosystem processes that characterize such forest ecosystems. The *Reassembly* Research Unit aims to understand the stability, in terms of resistance and resilience, of a tropical forest ecosystem and many of its components – based on space-for-time substitution of community and network recovery, underlying trait rules, and consequences for ecosystem processes (Figure 1). While *Reassembly* primarily targets fundamental research questions on network reassembly, its results inform nature conservation and policy makers on the recovery potential of tropical forests, potentially improving the restoration of tropical forests (Tylianakis et al., 2010; Pocock et al., 2012; Kaiser-Bunbury et al., 2015, 2017).

### 2.1 Study Site

Our study is located within the Ecuadorian Chocó, in the lowland rainforest of the Canandé watershed, near the town of Hoja Blanca in the Esmeraldas Province. The climate is typical for equatorial lowland rainforest, with mean annual temperature of ca. 22–23°C and mean annual precipitation of 3000–6000 mm with a dry season between June-July and October-November (based on the nearest weather stations 20–50 km away: Santo Domingo: 2800 mm, Luis Vargas Torres: 6200 mm). On-site weather data collection is currently undergoing with rain gauges and data loggers, and we expect to update this information in the future. Current land use in the area is relatively recent, occurring within the last 50 yrs according to local residents. A large portion of the old-growth forest in the area has been logged by timber industries and transformed by local people into cacao plantations, and less commonly into oil palms, as well as pastures for cows and horses to sustain their livelihood. Many plantations and pastures have been abandoned at different times, often to be included in private rainforest reserves, leaving a mosaic of habitats that differ in their time of recovery. The private reserves *Canandé* and *Tesoro Escondido* and surroundings (0.5°N 79.2°W, 130–540 masl) are 14,000 and 1,800 ha in size, respectively, and *Canandé* is still growing. Both harbor different habitats that range from old-growth forests to active plantations and pastures, including secondary forests of different regeneration ages. *Fundación Jocotoco* established *Canandé* Reserve in 2000 and immediately included pastures and plantations in their protected area. The *Tesoro Escondido* Reserve was established in 2016, with around 90% of primary forest, the rest being pastures of different ages and old cacao plantations. These active and old pastures and cacao plantations provide the basis for the chronosequence sites in this project.

We established 62 plots of 50 x 50 m (0.25 ha) along a chronosequence of forest regeneration in *Canandé* and *Tesoro Escondido* reserves during 2021–2022 (Figure 2a). Larger plots were difficult to implement owing to the topography of the area, which is hilly with many streams. The large number of plots implemented increases the spatial resolution of the chronosequence allowing well-replicated spatial data. We included cacao plantations and pastures as the main land-use types in the area to understand whether different land-use legacies lead to differences in resilience (Guariguata & Ostertag, 2000; Jakovac et al., 2021). Six plots are located in active cacao plantations, six in active pastures, 17 in regenerating cacao plantations, 16 in regenerating pastures, and 17 in old-growth forests (Figure 2a; Table 1).

**Figure 2.**
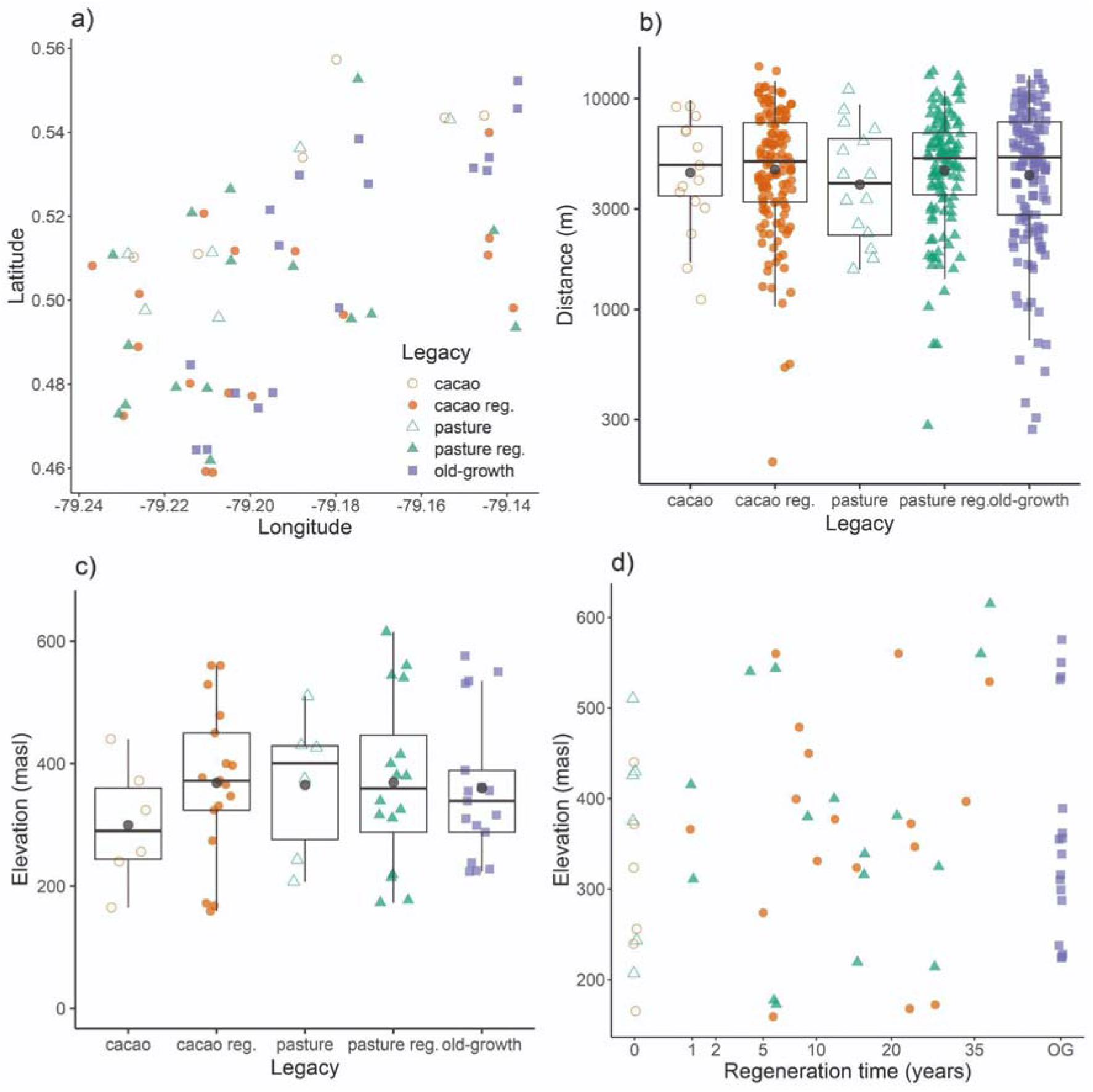
Plot spatial characteristics. a) Location of the 62 study plots at the Canandé and Tesoro Escondido reserves in the lowland rainforests of northwestern Ecuador. b) Distances between plots within each legacy type. The Y axis is log-transformed to facilitate the visualization of low values. c) Elevational distribution of each legacy type. d) Elevation is not correlated with square-root transformed regeneration time in all active and regenerating cacao and pasture plots. Old-growth forest (OG) plots were not included in the analyses because their time without human disturbance is unknown. Dark circles in all boxplots represent mean values.

**Table 1.** Mean values and standard errors of plot characteristics and response variables. Maximum tree height was calculated as the average of the five tallest stems per plot while maximum tree DBH was calculated as the average of the five widest stems per plot. Maximum tree height, maximum tree DBH, basal area, and number of stems were estimated for wild trees with DBH ≥ 7.95 cm. Reversed TSF was calculated as 1-TSF. Aboveground biomass (AGB) was estimated using species-specific wood density along with DBH and height data, and rarefied tree species richness was estimated with a coverage of 0.75 for wild trees with DBH ≥ 7.95 cm.

| Variable | Cacao | Regenerating cacao | Pasture | Regeneration pasture | Old-growth |
| --- | --- | --- | --- | --- | --- |
| Number of plots | 6 | 17 | 6 | 16 | 17 |
| Elevation (masl) | 299.5 ± 40.46 | 368.52 ± 30.63 | 365.16 ± 47.93 | 369.31 ± 34.82 | 360.05 ± 28.84 |
| Regeneration time (yrs) | 0 | 15.82 ± 2.61 | 0 | 15.12 ± 2.97 | NA |
| Maximum tree height (m) | 10.44 ± 2.39 | 24.03 ± 1.3 | 15.48 ± 2.24 | 21.82 ± 2.72 | 35.54 ± 0.92 |
| Maximum tree DBH (cm) | 19.2 ± 5.53 | 46.13 ± 3.34 | 22.1 ± 4.05 | 38.7 ± 4.36 | 93.99 ± 4.9 |
| Basal area (m<sup>2</sup>/ha) | 1.12 ± 0.46 | 14.29 ± 1.87 | 1.24 ± 0.39 | 12.27 ± 2.21 | 39.54 ± 2.58 |
| Number of stems | 6.67 ± 1.67 | 119.47 ± 10.41 | 9.33 ± 2.46 | 105.5 ± 16.9 | 178.35 ± 7.32 |
| Vertical vegetation heterogeneity | 1.07 ± 0.14 | 1.64 ± 0.04 | 0.47 ± 0.2 | 1.49 ± 0.08 | 1.86 ± 0.01 |
| Reversed TSF (%) | 61.79 ± 8.71 | 78.16 ± 1.43 | 53.18 ± 9.6 | 74.68 ± 2.82 | 82.29 ± 1.38 |
| AGB | 6.35 ± 3.55 | 82.11 ± 13.62 | 7.7 ± 3.35 | 67.42 ± 14.63 | 358.54 ± 30.72 |
| Tree species richness | 4.5 ± 1.99 | 24.25 ± 4.4 | 5.27 ± 1.56 | 20.63 ± 4.54 | 71.8 ± 6.45 |

A correct estimation of the regeneration time for the plots in our chronosequence was fundamental for a proper assessment of forest recovery. Regeneration time was determined as the number of years that have passed since abandonment until observational data were finished in 2023. Year of abandonment of the plots was established based on the documentation by *Fundación Jocotoco* when land purchasing began. Before this, the year was estimated based on interviews with previous landowners and park rangers. In addition, we refined these data using satellite-based (see Landcover context section). Regenerating cacao and pasture plots have been under natural regeneration without major human disturbance for 1–38 yrs (Table 1; Table S1). The mean (± SE) time of regeneration for cacao (15.8 ± 2.6 yrs) and pasture plots (15.1 ± 3 yrs) is similar (*t* = 0.18, *p* = 0.86). Active cacao plantations and pastures belong to farms located around regenerating plots and were assigned a regeneration time of zero because they are currently under use and represent typical conditions at which regeneration may start. Old-growth forests were not assigned a regeneration time because there was minimal evidence of human disturbance.

Active cacao plantations are monocultures of *Theobroma cacao* L. (varieties ‘National’ and ‘CCN51’) usually below 5 m in height, however, trees can reach up to 10 m in older plantations. Cacao trees, planted from seeds in the study area, are spaced out by 2–4 m each, often as part of mixed polyculture including coffee or fruit trees. Plantations generally lack shade trees, and herbicide application is common. Active pastures are mostly grazed by cattle and occasionally by horses. They consist mainly of pasture grasses such as *Brachiaria* (Trin.) Griseb. or *Axonopus scoparius* (Flüggé) Kuhlm., but some single large remnant shade trees or palms have been left. Smaller tree islands can be found along creeks. Old-growth forests contain large slow-growing trees of potential timber use and showed no signs of tree harvesting such as tree stumps or skid trails seen elsewhere. Besides, several species with high wood density values that are usual members of old-growth forests such as *Licania* spp., *Pouteria* spp., *Eschweilera* spp., or *Lecythis* spp. have been recorded in the old-growth forest plots.

### 2.2 Suitability of the spatial design

We aimed to select plots suitable for statistical analysis of forest recovery based on regeneration time and land-use legacy (cacao or pasture). Importantly, we selected plots from potential candidates that increased the spatial independence (e.g., large distances between plots of the same category and similar age) while avoiding an elevation bias or other confounding variables like amount of forest around the plots or land-use duration. The minimum distance between plots of the same type was 184 m between active cacao plots (Figure 2b). For the other legacies, the minimum distance between two plots was over 250 m. Mean distances between plots were 5.2 ± 1.1 km for active cacao plantations, 5.5 ± 0.7 km for regenerating cacao plots, 4.6 ± 1.1 km for active pastures, 5.3 ± 0.7 km for regenerating pastures, and 5.5 ± 0.77 km for old-growth forest plots.

Elevation bias within different reachable parts of the study area was reduced to the best extent possible within the limitations of the land-use history. Elevation (altitude in m above sea level) in our study plots ranges between 159–615 masl (Figure 2c; Table 1; Table S1). Mean elevation in active and regenerating cacao plots (350.5 ± 25.4 masl) was similar to active and regenerating pasture plots (368.2 ± 27.9 masl; *t* = −0.47, *p* = 0.64). Mean elevation in old-growth forests (360 ± 28.8 masl) was also similar to active and regenerating cacao plots (*t* = −0.25, *p* = 0.8) and to active and regenerating pastures (*t* = 0.2, *p* = 0.84). There was no correlation between elevation and (square-root transformed) regeneration time across all the active and regenerating cacao and pasture plots (Pearson’s *r* = 0.17, *p* = 0.26; Figure 2d), and neither for cacao (Pearson’s *r* = 0.22, *p* = 0.31) nor for pasture alone (Pearson’s *r* = 0.13, *p* = 0.56). Note that square-root transformation of regeneration time is a common method for obtaining annual growth rates because it approximately linearizes the slope (Hoenle et al., 2022).

The estimated duration of land use, either as pasture or cacao plantation, of the plots before abandonment was determined also through interviews, and by examining historic Landsat data (see Landcover context section below). We acknowledge that the accuracy of these estimates are low for many plots. Land-use duration of all active plots ranges between 6–23 yrs with a mean of 15.5 ± 2.1 yrs (Table S1). Regenerating plots had a larger range of duration use (1–30 yrs; 11.4 ± 1.2). There were no differences in duration use between cacao and pasture plots (*t* = −0.59–0.09, *p* = 0.56–0.93).

Even though there was no bias in elevation and land-use, analyses along chronosequences benefit from the consideration of biotic and abiotic environmental variation as potential explanatory variables besides the time of regeneration, including variables that are independent of recovery. The lack of such variables has led to criticism of chronosequence approaches (Johnson & Miyanishi, 2008; Elsy et al., 2023; but see Walker et al., 2010). Our abiotic baseline data across our chronosequence included temperature, humidity, soil composition, canopy cover, precipitation, and landscape composition derived from satellite data (Table S1).

### 2.3 Landcover context

To understand landscape effects across the chronosequence plots, we developed a landcover map (Figure S1) for active-to-regenerating agriculture and forest within 1 km of each study plot, a scale relevant for many organisms including invertebrates to smaller vertebrates. Obtaining precise maps over time is a challenge in lowland rainforests dominated by high cloud cover. Thus, we used an integrated approach combining different satellite imagery sources, ground in-situ verification, and local knowledge. Boundaries of current and historic land use were manually digitized from combined sources in ArcGis (Esri, 2023). As the primary base layer, we used 0.5 m resolution optical imagery with an accuracy of 5 m from World Imagery with cloud-free images for 22 Jan 2017 (north, 12 plots) and 13 Nov 2020 (south, 50 plots; Esri 2023). Shrub-successional and canopy-closure phases ≤ 15 yrs were generally visible on optical imagery with differences between early and late regeneration. To quantify the borders of older regeneration, we used time-series analysis of historic Landsat images from the European Commission’s Joint Research Centre for change in forest cover of Tropical Moist Forest (JRC TMF) which attempts to quantify the year when forest degradation occurred since 1990 (Vancutsem et al., 2021), as well as Global Forest Watch (GFW) analysis of changes in forest height since 2000 (Hansen et al., 2013). Remote sensing data were ground-truthed in the field by mapping changes in forest structure and composition along trails (e.g., areas dominated by *Cecropia* spp.). For regenerating cacao and the oldest regenerating pasture plots which were not visible on optical imagery or historic Landsat, we quantified approximate disturbance areas in the field. Fieldwork was conducted with original settlers or their families who had lived on the land and knew the timing of agricultural abandonment. Local knowledge and structural changes observed on the ground matched historic Landsat ∼2000 when extensive forest clearing occurred after the completion of the main logging road. Landsat maps also supported field data and local knowledge indicating minimal disturbance at old-growth plots. However, without complete Lidar coverage it was not possible to identify disturbances > 20-30 yrs including selective logging known to have occurred in easily accessible areas < 1 km from the main road since the 1960s (e.g. Endesa Botrosa concession). Thus, current forest cover maps represent areas dominated by mature trees but do not differentiate changes in forest structure associated with old-growth.

We quantified the location of *Reassembly* plots in a mosaic of agricultural land uses and forest including distance to the forest edge, patch size, and forest cover at three spatial scales, defined as the percentage classified as forest divided by the total area within three different radii (1 km, 500 m and 100 m; Table S2). Plots were located in a relatively intact landscape and overall forest cover averaged 74 ± 2.8 % within a 1 km radius, including regenerating forest. Most plots were near forest, and on average cacao and pasture plots were located 58 ± 7 m from the nearest forest edge within 10.1 ± 2.1 ha patches except for regenerating cacao plots which were in smaller 2.0 ± 0.4 ha cacao plantations (Figure 3a).

**Figure 3.**
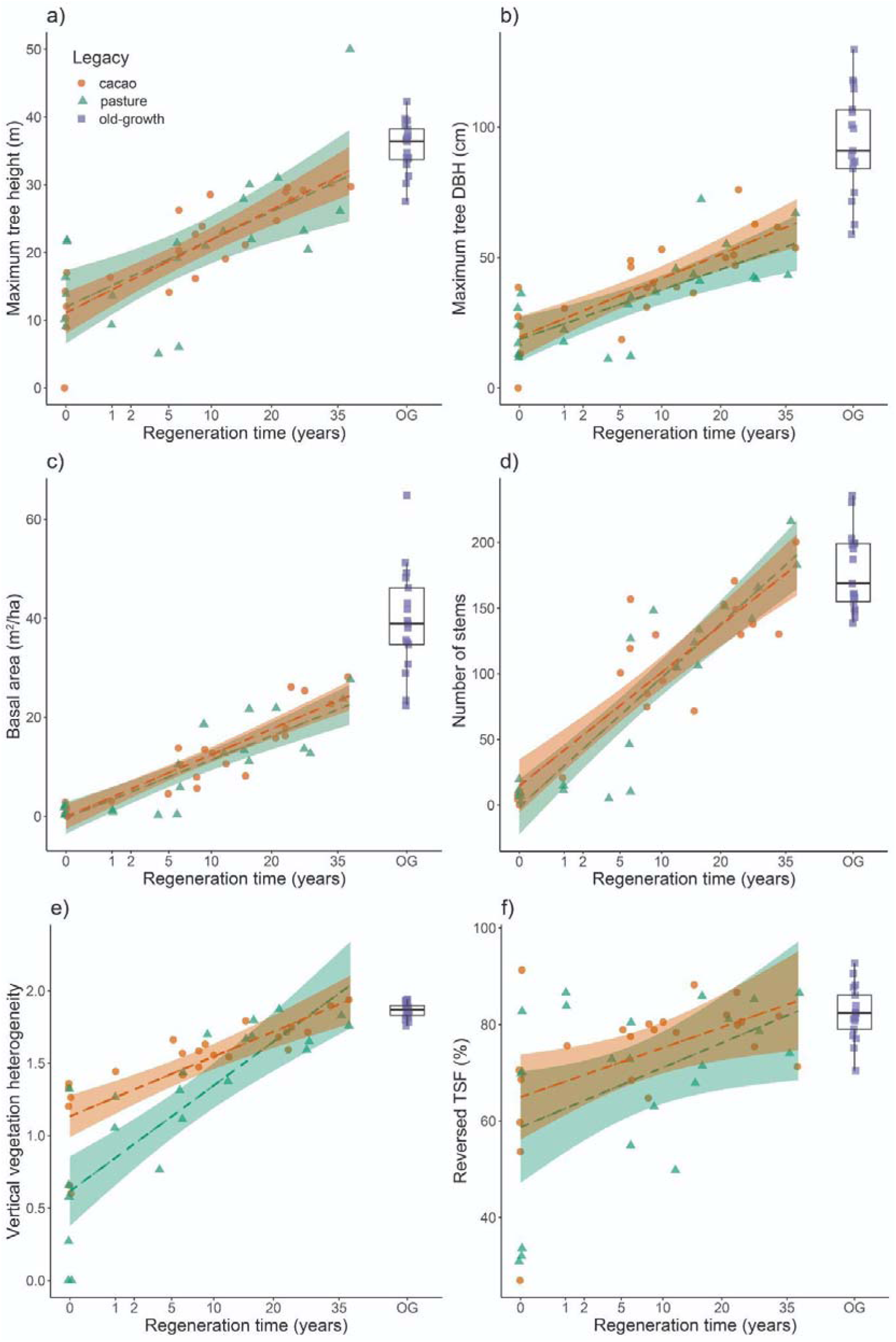
Recovery of forest structure. a) Maximum tree height, b) maximum tree DBH, c) basal area, d) number of stems, e) vertical vegetation heterogeneity, and f) reversed total site factor (TSF) increase significantly with regeneration time in cacao and pasture plots. Maximum tree height was calculated as the average of the five tallest stems per plot while maximum tree DBH was calculated as the average of the five widest stems per plot. Maximum tree height, maximum tree DBH, basal area, and number of stems were estimated for wild trees with DBH ≥ 7.95 cm. Reversed TSF was calculated as 1-TSF. Old-growth forest (OG) plots were not included in the analyses because their time without human intervention is unknown and were used only as a reference.

Old-growth plots averaged 388 ± 72 m to the nearest forest edge and at the larger spatial scale (within a 1 km radius), were surrounded by 87 ± 4 % forest, an 18% increase compared to agricultural plots at 69 ± 3 % (*F* = 10.6, *p* = 0.001). (Figure S2a, S2b). At smaller spatial scales (within a 100 m radius), old-growth forest plots were exclusively surrounded by 98 ± 2 % forest whereas forest cover was minimal around cacao and pasture plots (active: 11 ± 1%, regenerating: 36 ± 1 %). There was no significant correlation between elevation and land cover within 100 m (Pearson’s *r* = 0.11, *p* = 0.37; Figure S2c; Table S3). Because large tracts of intact forest tend to be retained in inaccessible areas such as ridgetops, there were weak significant correlations between elevation and landcover at 500 m and 1 km scales (Pearson’s *r* = 0.26–0.43, *p* = 0.0003–0.03) (Figure S2d, S2e). Additionally, land cover at different scales was significantly correlated with (square-root transformed) regeneration time (Pearson’s *r* = 0.38–0.8, *p* = 1.583e-15–0.002; Figure S2f, S2g, S2h; Table S2).

## 3. Recovery of forest structure, AGB, and tree species richness

Trees are key components of tropical ecosystems because they provide the structure and microclimate of the forest where animals can coexist and also form the base for key ecological interactions such as pollination, seed dispersal, and herbivory. We therefore estimate the time for total recovery, and the stability in terms of resistance and resilience of forest structure variables, aboveground biomass (AGB), and tree species richness in the 62 plots. We included as forest structure variables maximum tree height, maximum tree diameter at breast height (DBH), basal area, number of stems, vertical vegetation heterogeneity, and light availability. For analysis, AGB was estimated using species-specific wood density, while tree species richness was estimated using rarefaction with a representative coverage level of 0.75. We run these calculations for wild trees with DBH ≥ 7.95 cm.

### 3.1 Methods

#### 3.1.1 Tree survey and identification

Our assessment of trees on each plot occurred from February 2022 to July 2023 and comprised tree species identification and labeling of all individuals ≥ 25 cm of circumference at 1.3 m above the ground (≥ 7.95 cm diameter at breast height, DBH), including palms and lianas. We also measured the height of each tree using a laser rangefinder/hypsometer (Forestry Pro II, Nikon). Given that the terrain in the area is irregular and some height measures could be overestimated, any tree > 50 m was limited to this height. If a tree had more than one stem ≥ 25 cm circumference, we counted and measured up to the four thickest stems. A botanical collection and silica samples for further genetic and chemical analyses were obtained from each tree species. Tree identification was performed at the Herbario Nacional del Ecuador–INABIO using the collections deposited there as references.

A total 7921 stems of 7494 cultivated and non-cultivated trees were surveyed, representing the actual forest structure,. Among these, we identified 859 tree species and morphospecies, with 39% of them identified up to species level and 25 % to genus level. In total, 1393 trees were identified to morphospecies level because they did not present leaves during the survey or because additional work at the herbarium is required. We built a second dataset of only wild trees, including remnant ones in active agricultural plots. We used this second dataset for statistical analyses, by removing 779 stems of 639 cultivated trees of 11 species, including cacao, coffee, coconut, and lemon trees common in the area. Finally, we built a third dataset considering only wild trees with DBH ≥ 10 cm to make our results on AGB and species richness comparable with others. In this dataset, 5393 stems of 5216 trees, and 733 wild tree species and morphospecies remained.

#### 3.1.2 Forest structure

The forest structure variables analysed using the second dataset were maximum tree height (average of the five tallest stems), maximum tree DBH (average of the five widest stems), basal area, number of stems, vertical vegetation heterogeneity, and light availability. We multiplied basal area results by 4 to provide estimated data per 1-ha. Here, light availability is measured as the total site factor (TSF; i.e., the total solar radiation including direct and diffuse light, relative to open conditions) at a height of 2 m at three random selected locations within each plot using hemispherical photography (Solariscope SOL 300B). The instrument measures the shading of forest canopies relative to open field conditions. Accounting for the theoretical path of the sun (depending on geographical orientation and latitude), it obtains an accurate sub-canopy light estimate (Canham et al., 1990). Light measurements were taken at the four corners of a 10 ⍰ 10 m square centered around the plot center. For analysis, we averaged these measurements per plot. We also reversed TSF values (1-TSF) to understand it as the change of canopy shading due to an increased appearance of leaves and stems along the chronosequence. To assess vertical vegetation heterogeneity, we estimated vegetation cover at heights of 0.5, 1, 2, 4, 8, 16, 32, and 64 meters within each plot. To do so, we established five circular sampling areas with a 20-meter radius per plot, one at the center and four others at the corners. For each circle, the Shannon-Wiener diversity index was calculated, and the mean value of these indices from all five circles was then averaged to determine the overall vertical vegetation heterogeneity for each plot (Bibby et al. 2000).

#### 3.1.3 Aboveground biomass (AGB)

For analysis, AGB was computed for the second dataset with wild trees with DBH ≥ 7.95 cm. We calculated AGB for each individual using species-specific wood density along with DBH and height data with the Chave et al. (2014) equation #4 in the package BIOMASS (Réjou-Mechain et al., 2017) and then added per plot. If the species identity was not available for different reasons, BIOMASS obtains a wood density value for the plant genus or the family, because wood density is phylogenetically conserved (Chave et al., 2005). For those stems with no identification to family level, BIOMASS obtains a mean wood density value per plot and uses it in the AGB calculations. We also provide as supplementary material a measure of AGB calculated with a different equation from Chave et al. (2005) using DBH, height, and assuming a constant wood density of 0.5 g/cm3 (Table S4). In Table S4 we also included AGB calculated for the first (all trees with DBH ≥ 7.95 cm) and for the third datasets (wild trees with DBH ≥ 10 cm). In addition, we calculated aboveground carbon (AGC) stocks per plot for the three datasets in BIOMASS without accounting for potential errors in height and DBH because that data was provided and not estimated (Table S4). We multiplied AGB and AGC results by 4 to provide data for 1-ha.

#### 3.1.3 Tree species richness

As with AGB, we used estimations of tree species richness for wild trees with DBH ≥ 7.95 cm for analyses. We estimated coverage-based rarefied species richness using the package iNEXT (Hsieh et al., 2016) based on a representative coverage level of 0.75. In addition, we provide raw species richness obtained with vegan (Oksanen et al., 2022) as supplementary material (Table S4).

#### 3.1.4 Statistical analysis

First, we performed linear models to determine how and by which factors the six measures of forest structure (maximum tree height, maximum tree DBH, basal area, number of stems, vertical vegetation heterogeneity, and reversed TSF), AGB, and tree species richness change along the chronosequence. Elevation, the percentage of forest at 100 m around the plots, and land-use legacy (cacao or pasture) interacting with (square-root transformed) regeneration time, and duration use were set as fixed effects. The interaction between legacy and (square-root transformed) regeneration time was included because we wanted to determine whether former cacao and pasture plots can regenerate at a similar time. Old-growth forests were not included in the models because of their unknown age and extent of their last natural or human disturbance (if any). We performed an ANOVA test on the resulting models to look for type II errors. For model validation, we first checked visually that the residuals of the models follow a normal distribution. To test the lack of variance homogeneity, we performed a Bartlett test and examined the Pearson correlation between fitted and residual values of the models. We determined the lack of spatial autocorrelation of the model residuals with a Moran’s I test from the package ape (Paradis & Schliep et al., 2019).

Second, we estimated the time for total recovery of the response variables for all active and regenerating plots and for each legacy type (cacao and pasture) based on a simplified linear model without elevation, the interaction between legacy and (square-root transformed) regeneration time, the percentage of forest 100 m around the plots, and duration of use. We defined the following linear trend for the response variable (*y*) over time (*t*):

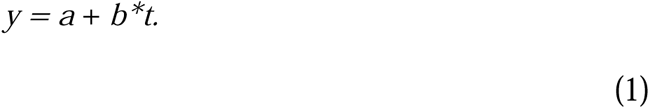

Here, *a* is the starting point (intercept) at time 0 and *b* is the slope of (square-root transformed) regeneration time. We used the median value of each response variable in old-growth forests (OG_median_) as reference to reveal the estimated recovery time as

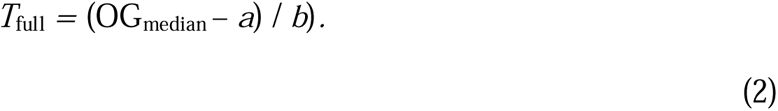

We then back-transformed (^2) the time for total recovery because we used square-root transformed regeneration time in the models.

Third, we calculated the resistance and resilience of the six variables of forest structure, AGB, and tree species richness across the regeneration gradient. Resistance and resilience were determined for cacao and pasture plots together and separately. Resistance was calculated as

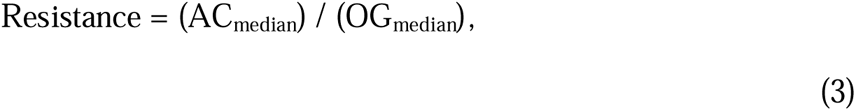

Where AC_median_ represents the median value of the response variables in active plots when regeneration time (*t*) is zero. Resilience was determined as a percentage/year using the following formula:

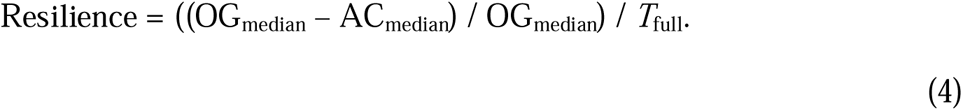

### 3.2 Results

Mean values of the forest structure variables in old-growth forests were always higher compared to those in regenerating and active plots (Table 1; Table S4). For instance, maximum tree height in old-growth forests was 35.54 ± 0.92 m while in regenerating plots it was 22.96 ± 1.47 m and in active plots it was 12.96 ± 1.74 m. Maximum tree DBH was 93.99 ± 4.9 cm in old-growth forests, 42.53 ± 2.76 cm for regenerating plots and 20.65 ± 3.3 cm for active plots. Basal area peaked at old-growth forests with a mean of 39.54 ± 2.58 m^2^/ha, while it was 13.31 ± 1.43 m^2^/ha and 1.18 ± 0.29 m^2^/ha in regenerating and active plots, respectively. Number of stems varied the most with an average of 178.35 ± 7.32 stems in old-growth forests, 112.7 ± 9.71 stems in regenerating plots and 8 ± 1.47 stems in active plots. Vertical vegetation heterogeneity in old-growth forests had a mean value of 1.86 ± 0.01, while in regenerating and active plots it was 1.57 ± 0.04 and 0.77 ± 0.15, respectively. Reversed TSF was 82.29 ± 1.38 % in old-growth forests, 76.47 ± 1.56 % in regenerating plots, and 57.49 ± 6.31 % in active plots.

The six forest structure variables increased significantly with (square-root transformed) regeneration time (Figure 3; Table S5). Vertical vegetation heterogeneity was influenced by land-use legacy and by its interaction with (square-root transformed) regeneration time (Figure 3e; Table S5). Neither elevation nor the percentage of forest at 100 m around the plots nor duration use influenced the change of the response variables along the chronosequence. Assumptions of the models including normal distribution of residuals and variance homogeneity were met except for vertical vegetation heterogeneity (Figure S3, Supplementary Results). Spatial autocorrelation was determined only for maximum tree DBH and vertical vegetation heterogeneity (Figure S3, Supplementary Results). Number of stems, vertical vegetation heterogeneity and reversed TSF were the parameters that recovered fastest at around 30 yrs, whereas maximum tree DBH was slowest to recover at 120 yrs (Table 2).

**Table 2.** Summary of total recovery (in years), resistance (%), and resilience (%/y) of forest structure variables, aboveground biomass (AGB), and tree species richness. maximum tree height was calculated as the average of the five tallest stems per plot while maximum tree DBH was calculated as the average of the five widest stems per plot. Forest structure variables, AGB estimated using species-specific wood density along with DBH and height data, and rarefied tree species richness estimated with a coverage of 0.75 were calculated for wild trees with DBH ≥ 7.95 cm. Reversed total site factor was calculated as 1-TSF.

| Variable | Total recovery (y) | Total recovery cacao (y) | Total recovery pasture (y) | Resistance (%) | Resistance cacao (%) | Resistance pasture (%) | Resilience (%/y) | Resilience cacao (%/y) | Resilience pasture (%/y) |
| --- | --- | --- | --- | --- | --- | --- | --- | --- | --- |
| Maximum tree height | 57.76 | 58.12 | 57.31 | 35.54 | 30.69 | 41.47 | 1.12 | 1.19 | 1.02 |
| Maximum tree DBH | 119.5 | 114.41 | 127.21 | 22.39 | 20.23 | 22.6 | 0.65 | 0.7 | 0.61 |
| Basal area | 103.88 | 101.84 | 106.82 | 2.52 | 2.4 | 2.93 | 0.94 | 0.96 | 0.91 |
| Number of stems | 30.82 | 29.83 | 32 | 4.44 | 4.44 | 4.73 | 3.1 | 3.2 | 2.98 |
| Vertical vegetation heterogeneity | 29.89 | 23.35 | 38.57 | 35.15 | 65.95 | 22.74 | 2.17 | 1.46 | 2 |
| Reversed TSF | 32.56 | 26.38 | 40.72 | 77.89 | 77.89 | 62.78 | 0.68 | 0.84 | 0.91 |
| AGB | 202.27 | 199.36 | 208.1 | 0.92 | 0.86 | 1.52 | 0.49 | 0.5 | 0.48 |
| Tree species richness | 101.2 | 99.09 | 104.24 | 6.27 | 4.3 | 7.04 | 0.93 | 0.97 | 0.89 |

Total above-ground biomass (AGB) calculated with species-specific wood density for all trees, including cultivated ones, was 2181.4 t within all the plots of the chronosequence, while AGB only of non-cultivated wild trees was 2163.55 t. This means that the AGB from cultivated trees in our chronosequence, mostly cacao trees, represents 17.85 t (0.8 %). When considering only wild trees with DBH ≥ 10 cm, total AGB was 2124.15 t. AGB was highly correlated with basal area (Pearson’s *r* = 0.97, *p* < 0.001). Within old-growth forests, the mean of AGB was 358.54 ± 30.72 t/ha (Table 1; Table S4) while in active plots it was 7.03 ± 2.34 t/ha and in regenerating plots it was 74.99 ± 9.9 t/ha. AGB increased significantly with (square-root transformed) regeneration time based on the ANOVA test (*p* < 0.001; Table 3). A normal distribution of residuals, variance homogeneity, and lack of spatial autocorrelation were determined (Figure S4, Supplementary Results). The model estimated a complete recovery of AGB at 202.6 yrs for all active and regenerating plots, 199.6 yrs when regenerating from cacao, and 208.6 yrs when regenerating from pasture (Table 2; Figure 4a).

**Figure 4.**
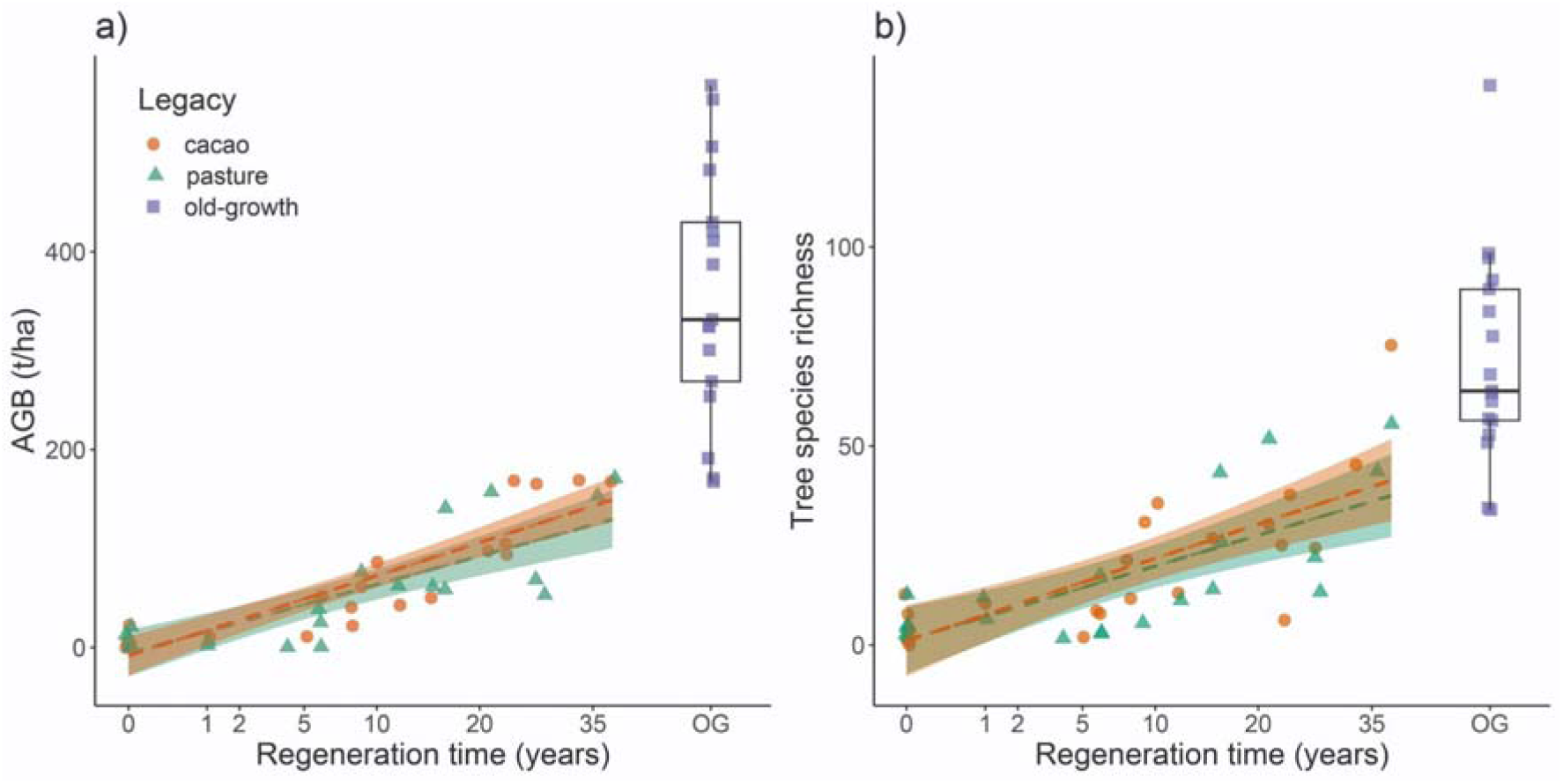
Recovery of aboveground biomass (AGB) and tree species richness. a) AGB estimated using species-specific wood density along with DBH and height data, and b) rarefied tree species richness estimated with a coverage of 0.75 increase significantly with regeneration time in cacao and pasture plots. AGB and rarefied tree species richness were calculated for wild trees with DBH ≥ 7.95 cm. Old-growth forest (OG) plots were not included in the analyses because their time without human intervention is unknown and were used only as a reference. Boxplots show the median value of the Y axis.

**Table 3.** Results of linear models. Aboveground biomass (AGB) was estimated using species-specific wood density along with DBH and height data, and rarefied tree species richness was estimated with a coverage of 0.75 for wild trees with DBH ≥ 7.95 cm. *P* values in bold < 0.05.

| AGB - ANOVA type II error |  |  |  |  |
| --- | --- | --- | --- | --- |
| Variable | Sum Sq | Df | F value | *p* |
| Elevation (masl) | 0 | 1 | 0.0003 | 0.987 |
| Forest 100 m around | 471 | 1 | 0.4947 | 0.4861 |
| Land-use legacy | 506 | 1 | 0.5318 | 0.4703 |
| sqrt(Regeneration time) | 75727 | 1 | 79.5234 | 7.46E-11 |
| Duration use | 256 | 1 | 0.2684 | 0.6074 |
| Land-use legacy (pasture) : sqrt(Regeneration time) | 426 | 1 | 0.4469 | 0.5079 |
| Residuals | 36186 | 38 |  |  |
| Tree species richness - ANOVA type II error |  |  |  |  |
| Elevation (masl) | 398.4 | 1 | 3.1849 | 0.08231 |
| Forest 100 m around | 175.2 | 1 | 1.4007 | 0.24396 |
| Land-use legacy | 29 | 1 | 0.2318 | 0.63294 |
| sqrt(Regeneration time) | 3507.9 | 1 | 28.0424 | 5.26E-06 |
| Duration use | 177.3 | 1 | 1.4175 | 0.24119 |
| Land-use legacy (pasture) : sqrt(Regeneration time) | 46.3 | 1 | 0.37 | 0.54664 |
| Residuals | 4753.5 | 38 |  |  |

Rarefied tree species richness reached an average of 71.8 ± 6.45 in old-growth forests (Table 1; Table S4), 22.5 ± 3.13 in regenerating plots and 4.88 ± 1.2 in active plots. Tree species richness also increased significantly only with (square-root transformed) regeneration time based on the ANOVA test (*p* < 0.001; Table 3). Assumptions of the model were also met (Figure S4, Supplementary Results). Complete recovery of tree species richness was estimated at 107.9 yrs for all active and regenerating plots, 104.9 yrs for cacao, and 112 yrs for pasture (Table 2; Figure 4b).

Resistance was generally slightly higher for pasture plots, potentially because of the remnant trees left standing in pastures, except for vertical vegetation heterogeneity and reversed TSF (Table 2; Figure 5). The highest resistance was detected for reversed TSF (77.89 %) while the lowest was for AGB (0.92 %). Resilience, on the other hand, was slightly higher for cacao plots except for vertical vegetation heterogeneity and reversed TSF (Table 2; Figure 5). The highest resilience was recorded for the number of stems (3.1 %/y) while the lowest was determined for AGB (0.49 %/y) too.

**Figure 5.**
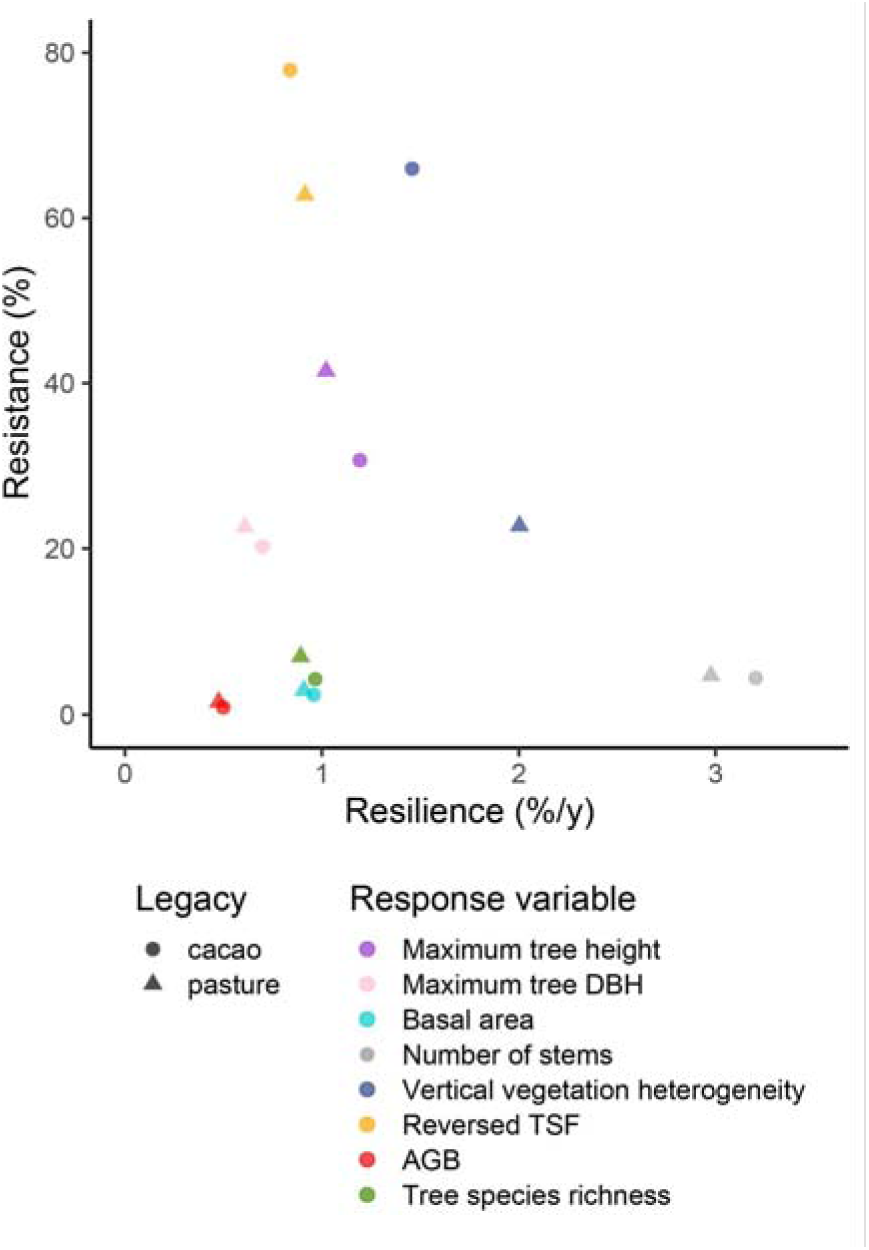
Recovery dynamics of forest structure, aboveground biomass (AGB), and tree species richness. Cacao and pasture plots (active and regenerating) are plotted separately. maximum tree height was calculated as the average of the five tallest stems per plot while maximum tree DBH was calculated as the average of the five widest stems per plot. Forest structure variables, AGB estimated using species-specific wood density along with DBH and height data, and rarefied tree species richness estimated with a coverage of 0.75 were calculated for wild trees with DBH ≥ 7.95 cm. Reversed total site factor (TSF) was calculated as 1-TSF.

### 3.3 Discussion

In an era of ongoing deforestation in the tropics with huge losses of biodiversity and ecosystem functioning, an important challenge today is to protect and restore such vulnerable ecosystems. This includes the urgent need to better understand drivers of successful natural recovery and community assembly of ecosystems, and natural disturbance–recovery dynamics. Chronosequences like the one presented in our study in the Ecuadorian Chocó have proven particularly useful to understand forest recovery over several decades, and to reveal successional patterns of various ecosystem attributes. One important goal in this endeavor is to distinguish resistance of properties to withstand disturbance and their resilience – both describing different dimensions of ecosystem stability. Here we presented the recovery of forest structure, aboveground biomass (AGB), and tree species richness, and showed how our spatial design of 62 plots was suitable to understand recovery trends over time without confounding effects of elevation, land-use duration, landscape composition, and spatial proximity. For example, we estimated a relatively slow recovery of tree richness (101 yrs), both due to low resistance (7%) and resilience (0.9 %/y). Studies addressing recovery within the Reassembly project will estimate resistance and resilience of various taxa, interactions and ecosystem components, providing a broader quantitative assessment of this complex rainforest system. For ants, Hoenle et al. (2022) determined that species richness recovered much faster (7-8 yrs), with high resistance (72%) and resilience (10.6 %/y), than community composition (21-29 yrs) across many of the plots of our chronosequence. This finding is consistent with other chronosequences elsewhere, where species diversity of animals generally recovered faster than their species composition (Curran et al., 2014). We predict that species interaction networks may recover even slower than species diversity and composition, particularly meta-community networks that have more complex features (Moreno-Mateos et al., 2020).

Overall, forest structure variables are less affected by deforestation than AGB and tree species richness based on the values of resistance. The forest structure variables directly related with tree size or density (maximum tree height and DBH, basal area, and number of stems) show relatively low resistance compared to variables indirectly related with trees such as vertical vegetation heterogeneity and reversed total site factor (TSF). These last two variables behave heterogeneously after disturbance among the active plots because not all plots have standing trees which influence vegetation complexity and the amount of light present. Particularly, reversed TSF showed the highest resistance suggesting that after forest disturbance there are still areas where light does not penetrate because of the remnant trees that could have been left standing or were planted during cultivation. Reversed TSF also shows relatively high resilience, which added to high resistance allows one of the shortest times for full recovery analyzed here. The low resistance and resilience for AGB results in the longest time for full recovery. The number of stems showed the highest resilience and also one of the shortest times for total recovery despite its low resistance, suggesting that recovery time is not limited by low resistance when resilience is high. Resistance and resilience for the variables analyzed here were in general similar for cacao and pasture plots, showing that land-use legacy does not influence recovery dynamics in our chronosequence as also revealed by our linear models.

The total recovery of the variables analyzed occurred at different times. The number of stems, vertical vegetation heterogeneity, and reversed TSF are the fastest variables to recover just after 30–32 yrs. On the other hand, maximum tree DBH and basal area recover after 120 and 104 yrs, respectively. Basal area recovers at a similar time compared to a tropical forest in Panamá whose recovery was estimated after 90 yrs (Elsy et al., 2023). This shows that despite regenerating forests having a similar number of stems compared to old-growth forests in a relatively short period of time, the full recovery of their structure still takes 4x that time. The full recovery of AGB estimated at 202 yrs takes almost 2x the time required for the recovery of forest structure potentially because of differences in species composition and wood density. In old-growth forests it is common to find tree species with high wood density compared to species found during early forest regeneration stages (Poorter et al., 2021a, b). This increases AGB in old-growth forests compared to regenerating forests with trees of the same size. Based on the variables analyzed here, the recovery of forest in Canandé would take at least 200 years. Nevertheless, the recovery of species composition and interaction networks is expected to take even a longer time (Crouzeilles et al., 2017; Moreno-Mateos et al., 2020).

## 4. Generalization to other chronosequences

A summary of many chronosequences from Neotropical forest sites has been published by the “2ndFOR” network (AGB from 41 sites: Poorter et al., 2016, tree species richness from 56 sites: Rozendaal et al., 2019, several forest attributes from 77 sites including 8 from West Africa: Poorter et al., 2021b). This allowed us to place our study design, environmental conditions, tree diversity, and recovery into a broader context.

Our chronosequence covers by far the largest total area of plots (4.25 ha of old-growth forests, 8.25 ha of regenerating forests, 3 ha of active agriculture) among all studies reviewed by Rozendaal et al. (2019) and thus involves a particularly large number of tree morphospecies (Figure 6a), even if the latter may become slightly lower once all trees have been identified to species. Hence, the chronosequence in the Chocó is particularly well resolved due to a large number of relatively large plots; only five other studies reviewed by Rozendaal et al. (2019) had more plots than ours but these were much smaller. Note, however, that most tree inventories used a lower threshold of DBH (5 cm) than ours (7.95 cm). Our study area is particularly wet (precipitation > 3000 mm) and its forest cover is relatively high (74%) compared to the majority of other chronosequences (Figure 6b). Our chronosequence is representative regarding its regeneration time range (0–38 yrs) compared to other neotropical studies (oldest plots: median 40 yrs, range: 15–100 yrs; Rozendaal et al., 2019).

**Figure 6.**
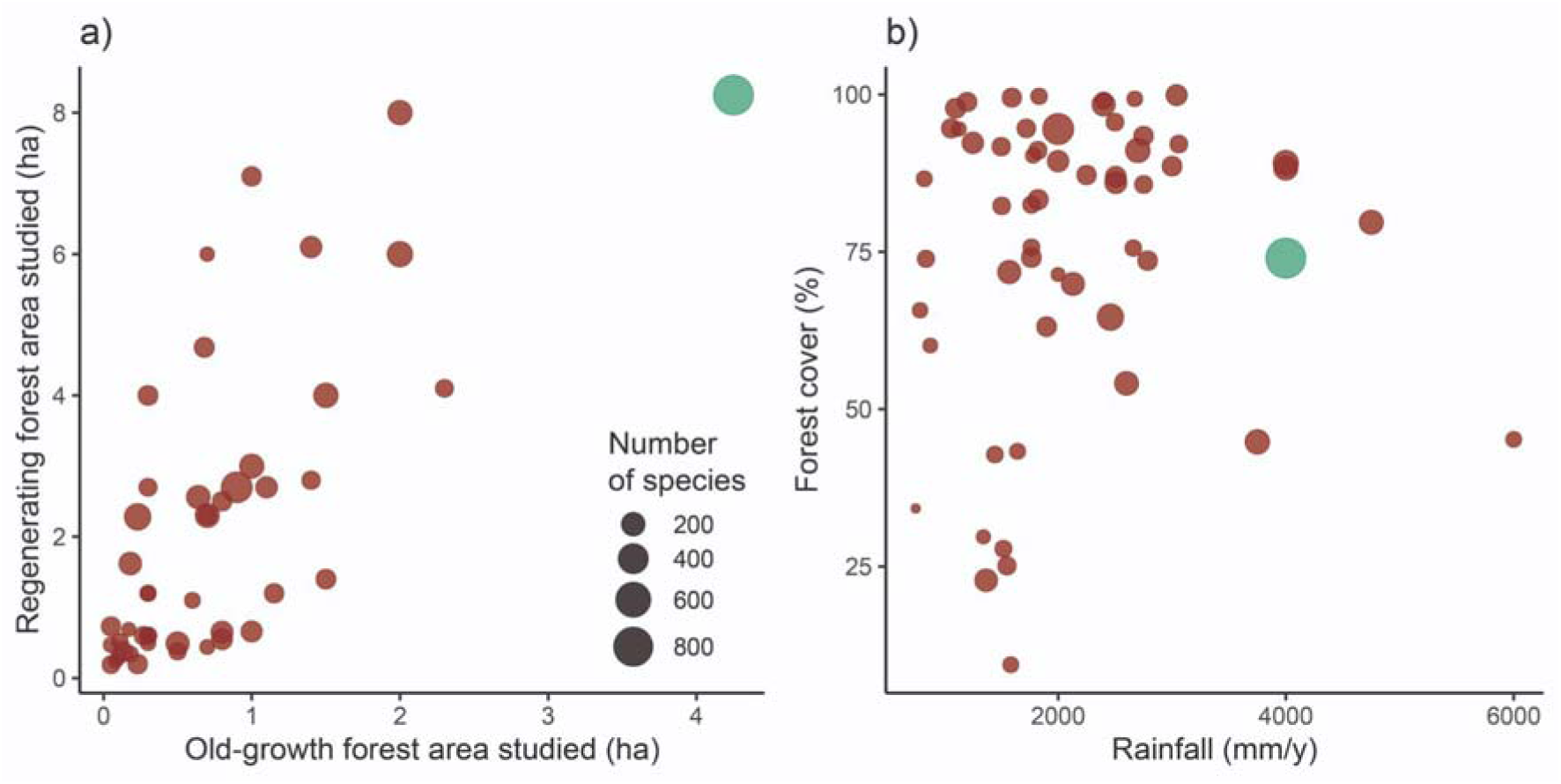
Context of our chronosequence (green) in 56 other forest chronosequences (red) from various dry to moist forests in the Neotropics (Rozendaal et al., 2019). Total area covered from all old-growth versus regenerating forest plots shown, as well as rainfall and forest cover in the landscape. Dots scaled by total number of tree morphospecies.

Aboveground biomass (AGB) in old-growth forests in our study (358 ± 31 t/ha, *n* = 16 plots, DBH ≥ 7.95 cm) is considerably higher than in many other Neotropical old-growth forest sites (mean 255 ± 117 t/ha, *n* = 19 studies mostly with DBH ≥ 5 cm; Poorter et al., 2016). The same was observed for our AGB estimates for larger trees (DBH ≥ 10 cm; mean AGB 354 ± 30 t/ha compared to other Neotropical forests calculated in the same way (mean of means 301 ± 31 t/ha, *n* = 14 sites, 146 1-ha plots, 333 0.1-ha plots; Poorter et al., 2015). The high precipitation in our chronosequence compared to others (Figure 6b) could explain the high AGB found because this variable is mostly driven by rainfall (Poorter et al., 2015). Beside AGB, aboveground carbon (AGC) for larger trees (DBH ≥ 10 cm) was 167 ± 14 t/ha. These were comparable to the values reported for a 1-ha permanent plot within the study area (AGB: 307.2 t/ha, AGC: 153.6 t/ha) for a total of 441 individual trees (DBH ≥ 10 cm) from 101 species (Lozano et al., 2023). AGB resilience in our study was similar to those in other Neotropical forests because we predicted an AGB of 123 ± 0.6 t/ha for 20 yrs secondary forests while the estimated average of 20 yrs plots across other neotropical chronosequences was 122 t/ha (range: 20–225□t/ha; Poorter et al., 2016).

The recovery of AGB in Canandé occurred over a much longer time compared to other chronosequences in other Neotropical forests. For instance, in a review of Neotropical secondary forests, it was determined that a median time of 66 yrs was enough to recover 90 % of AGB levels of old-growth forests (Poorter et al., 2016). In our case, it would take 164 years to recover 90 % of AGB, which is 2.5x the time estimated for the region. A meta-analysis across pairs of tropical secondary vs. old-growth forests revealed that complete recovery of AGB was reached after an average of 80 yrs (Martin et al., 2013), while we estimate this process would again take 2.5x longer in our system. These comparisons of AGB with other chronosequences may suggest that old-growth forests in Canandé have been mostly undisturbed during the last decades, accumulating biomass through a relatively long period of time.

The original datasets of tree diversities per plot and recovery times per study from the “2ndFOR” compilations were not available, hence we only compare the averages reported. Moreover, estimates for total recovery time differ strongly between statistical methods. The mean species richness per 25 tree stems (rarefaction method) was 17.9 ± 1.8 across the 16 old-growth plots in our study, higher than the 14.7 ± 4.5 found across 43 studies summarized in Rozendaal et al. (2019) using this rarefaction method. This difference in tree species richness could be explained by the larger forest area studied here (Figure 6a), allowing the finding of rare species that increase this diversity measure. Recovery time estimates in the meta-analysis of Poorter et al. (2021b) were obtained by fitting a saturating function, and total recovery was then defined for the predicted function reaching 90% of the old-growth level. Applying this method to our data results in a predicted recovery time of 55 years, well comparable with the mean of 54 years (11–228 yrs) reported by Poorter et al. (2021b). Overall, this shows that our highly resolved chronosequence was typical for other neotropical sites, being relatively rich in tree species and with a high biomass density, and with comparable recovery rates.

These comparisons against other chronosequences show that the natural regeneration of the forests in Canandé would take a longer time than for others in the region, taking even centuries. Although ecological restoration success is higher for natural regeneration than for active restoration (Crouzeilles et al., 2017), some assistance to natural regeneration could accelerate the recovery of variables such as AGB. For instance, trees from species with high wood density typical from old-growth forests (Poorter et al., 2021a, b) could be planted in regenerating forests that already have a structure that allows the survival of these planted trees. In that way, the recovery of AGB could happen over a shorter time span. On the other hand, variables that recover faster such as those related with forest structure or species diversity could be dependent only on the propagules arriving from the relatively high forested landscape around the plots (Figure 6b).

## 5. Outlook and future directions

Although deforestation by far outweighs the areas of recovering tropical forests, increasing efforts are being taken to protect and restore rainforest ecosystems from agricultural land or heavily exploited forests. The recovery of secondary forests is expected to play an increasing role in sustainable timber production, water protection, and biodiversity management during the next decades (Poorter et al., 2016; Chazdon & Guariguata, 2016). However, these efforts need to be implemented based on scientific grounds if the aim is to increase their chances of success (Brancalion et al., 2019). The implementation of chronosequences to study the recovery of ecosystem processes maintained by different taxonomic groups can promote an urgently needed understanding of the recovery potential of tropical forests, important components involved and thus contribute to successful restoration and conservation. Some first studies analyzing the reassembly of different animal communities and ecosystem pools such as deadwood in Canandé have been recently published (Hoenle et al., 2022; 2023; Müller et al., 2023; Falconí-López et al., in press).

Moving forward, we will test predictions of diversity and network reassembly and empirically assess the dynamics of species interaction networks along our chronosequence. Targeted networks and ecological processes include pollination, seed dispersal, seedling establishment, herbivory, predation, and dead wood decomposition, and thus multiple important mutualistic and antagonistic relationships between animals and plants. Different types of traits may enhance population and network resistance and resilience, respectively, and determine the rate of ecosystem processes to which these networks contribute (Sakschewski et al., 2016). In general, we expect that the diversity of plant and animal communities, species community composition, and species interaction networks resemble those of mature old-growth forests with time. However, different reassembly trajectories of species communities and interaction networks should be determined by evolutionary history and background taxonomic and functional diversity (Farneda et al., 2021; Molina-Venegas et al., 2018; Mahayani et al., 2020).

Whereas chronosequences have revealed many insights in changes in tree biomass, species’ abundances, diversity, and composition with time, data on species interactions or chronosequences looking at multiple taxa analyzed simultaneously remain scarce. Only a few studies have investigated the reassembly of interaction networks along chronosequences (see Staab et al., 2016; Redmond et al., 2019; Montoya-Pfeiffer et al., 2020), and thus a mechanistic understanding of the network dynamics is lacking. The degree of specialization of interaction networks varies across interaction types. Hence, mutualistic networks comprising pollinators, and primary and secondary seed dispersers are expected to be essential for forest recovery, particularly for maintaining the diversity of trees (Jordano, 2000; Ollerton et al., 2011). In turn, antagonistic interactions such as seed predation, seedling herbivory, or their reduction by predators, parasitoids, and pathogens, can also play an important role in the reassembly of tree communities (Bagchi et al., 2014). Our functional understanding of how communities and ecosystems reassemble, and why some systems return to a natural state while others do not, strongly depends on understanding these interaction networks.

## Acknowledgements

This work was supported by the Deutsche Forschungsgemeinschaft (DFG) funded Research Unit REASSEMBLY (FOR 5207). REASSEMBLY is a collaborative project between German and Ecuadorian institutions. We thank the *Fundación Jocotoco* and *Fundación Reserva Tesoro Escondido* for logistic support and permission to conduct research on their reserves. We also thank Katrin Krauth and Julio Carvajal for logistical assistance, Bryan Tamayo for plot managing, and Fredi Cedeño, Franklin Quintero, Jerson Loor, and Johan Párraga for field assistance during the tree survey. For landcover mapping in the field, we greatly appreciate the assistance and local knowledge of Ramón Vélez, Patricio Paredes, Alcides Zambrano, Amado De la Cruz, Lady Condoy, and Silvia Vélez. We would like to especially acknowledge Adriana Argoti, Leonardo de la Cruz, Jefferson Tacuri, and Ismael Castellanos for their help during plot establishment. We greatly thank Robin Chazdon for useful comments provided to this manuscript. We acknowledge the Ministerio del Ambiente, Agua y Transición Ecológica for granting collection and research permits under the Genetic Resources Access Agreement number “MAATE-DBI-CM-2021-0187”.

## Boxes

**Glossary - Concepts for temporal trends, disturbance, and stability**

### Chronosequence

A series of plots in a region, with comparable site attributes, that primarily differ in time span after a specific event (Bakker et al. 1996, Walker et al. 2010, Dunn 2004). The **chronosequence** in our study represents forest recovery (natural succession) from **disturbance** (agricultural use).

### Disassembly

An altered community composition and network characteristics immediately following a perturbation, typically affecting a fraction of the original set of species, relationships, or interactions.

### Disturbance

A relatively discrete event that disrupts components of an ecosystem, e.g. its community or population structure, resource or substrate availability and environmental conditions, and is more broadly defined as perturbation (Pickett & White 1985). May involve human influences as well as natural causes of fluctuations, e.g. forest gaps created by tree falls and small-scale mosaics of succession.

### Intermediate disturbance hypothesis (IDH)

A hypothesis that states that repeated disturbances at an intermediate level promote species coexistence via maintaining a non-equilibrium state, thus providing a release from competitive dominance of particular species (Connell 1978). Explicitly focusing on tropical rainforest trees, Connell suggested that “the highest diversity (…) should occur either at an intermediate stage in succession after a large disturbance or with smaller disturbances that are neither very frequent nor infrequent”. The IDH thus makes an explicit prediction for changes in tree richness (upper graph) and composition (lower graph) with forest recovery.

### Reassembly

The process of re-organization of community composition and network characteristics in an ecosystem that recovers from a perturbation.

### Recovery

The process of regaining original ecosystem properties and any measure after disturbance until levels of the reference state are reached (**recovery time**). Recovery includes reassembly that is more narrowly defined for more complex community composition or network properties.

### Resilience

One aspect of temporal **stability**: the "rebounding" of a system following a disturbance. While the concept of resilience may involve different capacities of reorganization to retain an equilibrium level or non-equilibrium states of a function, structure, identity or relationship (McCann 2000), its most applicable metric is simple: **recovery rate** (Pimm 1984). It could be defined as a rate of the initial loss (i.e. as the slope of the trajectory from the disturbed level to the reference level, or percent of losses regained in a given time) avoids that resilience and resistance are confounded (Pimm 1984, Pimm et al. 2019, White et al. 2020). Our project thus aims at quantifying resilience, and at studying *mechanisms* contributing to its variation.

### Resistance

Another aspect of **stability** of an ecosystem: the act of opposing and withstanding a disturbance or perturbation. Resistance is measured as the proportion of an initial property that remains following the disturbance (or while a disturbed phase continues), applicable to ecosystem characteristics such as species composition, network properties or ecosystem processes (Pimm 1984, Pimm et al. 2019, White et al. 2020).

