## Supplementary Material for "Reassembly of a tropical rainforest ecosystem: A new chronosequence in the Ecuadorian Chocó tested with the recovery of tree attributes"

^1^ Grupo de Investigación en Ecología y Evolución en los Trópicos-EETrop, Universidad de Las Américas, Quito, Ecuador.

^2^ Florida Museum of Natural History & Department of Biology, University of Florida, Gainesville, Florida, USA

^3^ Division of Conservation Biology, Institute of Ecology & Evolution, University of Bern, Bern, Switzerland.

^4^ Senckenberg Biodiversity and Climate Research Centre (SBiK-F), Frankfurt am Main, Germany.

^5^ Institute for Ecology, Evolution & Diversity, Goethe University Frankfurt, Frankfurt am Main, Germany.

^6^ Field Station Fabrikschleichach, Biocenter, University of Würzburg, Rauhenebrach, Germany

^7^ Ecological Networks Lab, Department of Biology, Technische Universität Darmstadt, Darmstadt, Germany
^8^ Fundación Jocotoco, Valladolid N24-414 y Luis Cordero, Quito, Ecuador.

* Corresponding author: Sebastián Escobar

**Supplementary Results**

Assumptions of the models for forests structure variables including normal distribution of residuals and variance homogeneity were met (all Pearson's *r* < 0.001, *p* = 1; Bartlett's *K*-squared = 0.007–1.603, *p* = 0.21–0.93; Figure S3), except for vertical vegetation heterogeneity (Bartlett's *K*-squared = 6.72, *p* = 0.009). No spatial autocorrelation of the models residuals occurred for maximum tree height, number of stems, basal area, and reversed TSF (Moran’s *I*_obs_ = -0.101–0.048, *I*_exp_ = -0.022, SD = 0.05, *p* = 0.12–0.96; Figure S3), but maximum tree DBH (Moran’s *I*_obs_ = -0.135, *I*_exp_ = -0.022, SD = 0.05, *p* = 0.03; Figure S3f) and vertical vegetation heterogeneity (Moran’s *I*_obs_ = 0.15, *I*_exp_ = -0.022, SD = 0.05, *p* = 8.69E-4; Figure S3o) did show spatial autocorrelation

The residuals of the model of AGB followed a normal distribution (Figure S4a), and variance homogeneity was discarded (Pearson's *r* < 0.001, *p* = 1; Figure S4b; Bartlett's *K*-squared = 0.03, *p* = 0.853). No spatial autocorrelation of the model residuals was detected (Moran’s *I*(obs) = -0.025, *I*(exp) = -0.022, SD = 0.052, *p* = 0.961; Figure S4c). Regarding, tree species richness, a normal distribution was observed in the residuals of the model (Figure S4d) and we discarded the presence of variance homogeneity (Pearson's *r* < 0.001, *p* = 1; Figure S4e; Bartlett's *K*-squared = 0.07, *p* = 0.791). Spatial autocorrelation was not detected (Moran’s *I*(obs) = 0.054, *I*(exp) = -0.022, SD = 0.051, *p* = 0.136; Figure S4f).

**Supplementary Figures**

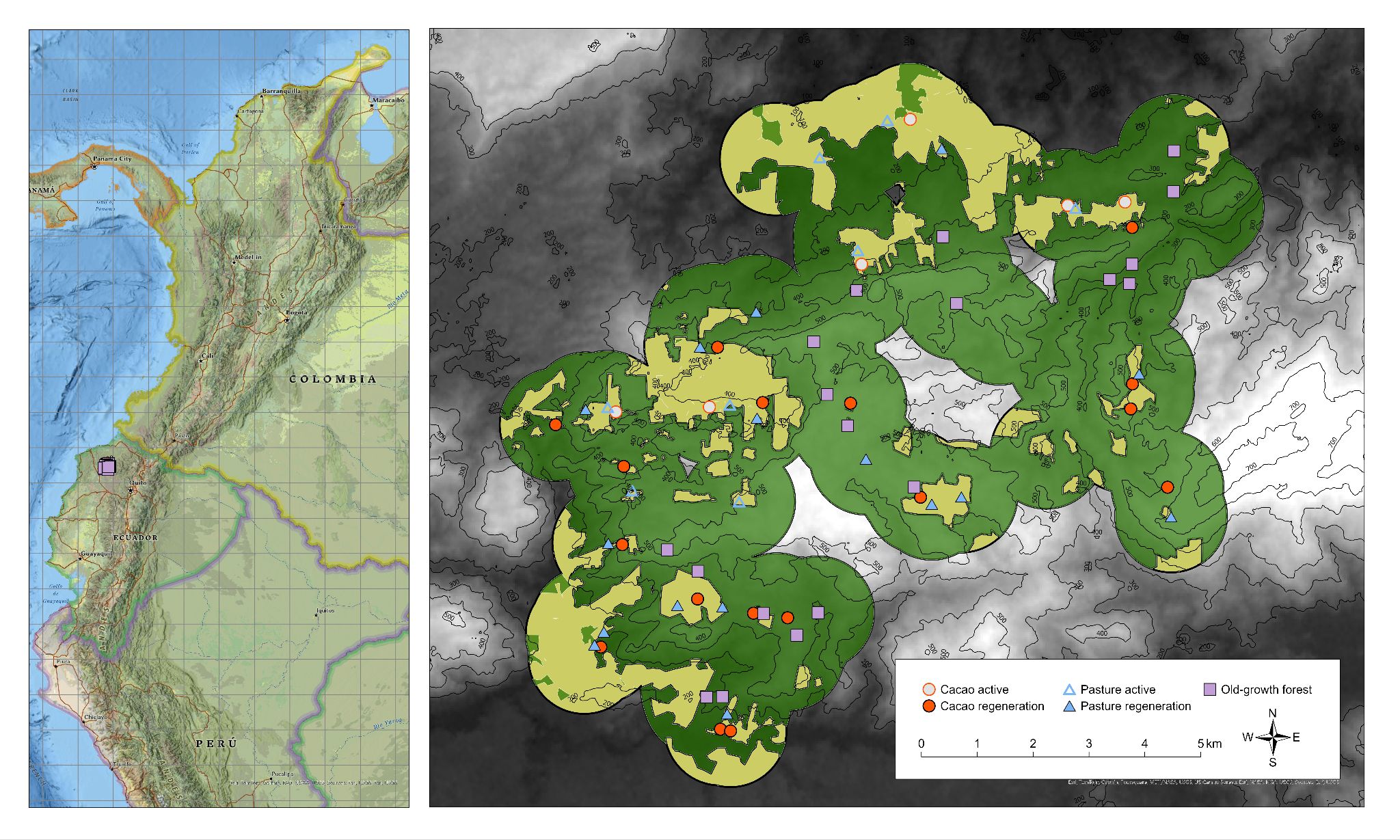

**Figure S1.** Distribution of the 62 study plots of the REASSEMBLY project in the Chocó

rainforest of Canandé, Ecuador. Symbols indicate different habitat and land-use types. Green indicates forest and tan indicates active/regenerating agriculture within 1-km of study plots.

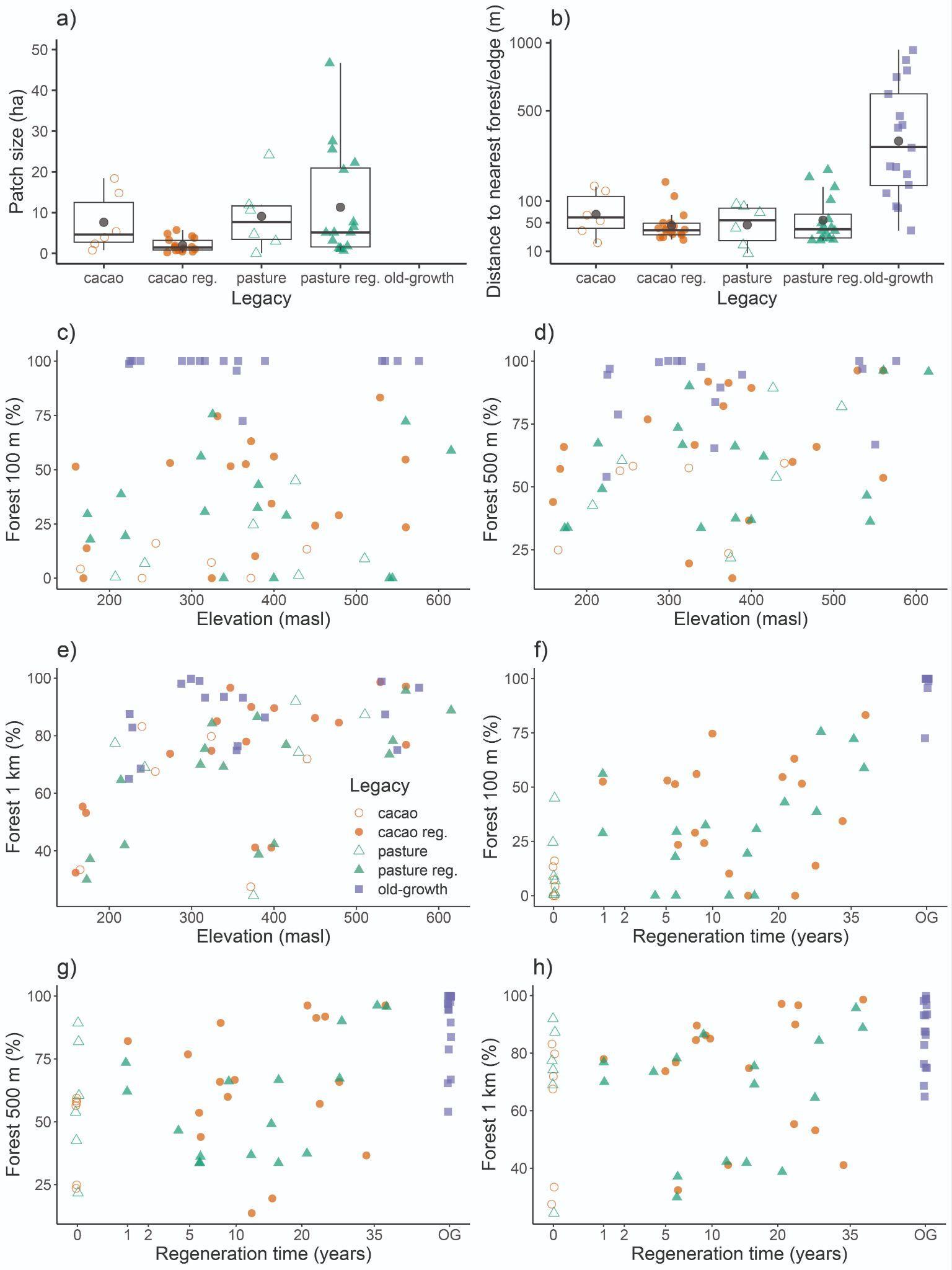

**Figure S2.** Plot landcover characteristics. a) Patch size (ha) within each legacy type. b) Distances to nearest forest (active and regenerating cacao and pasture plots) or nearest forest edge (old-growth forest plots). The Y axis is log-transformed to facilitate the visualization of low (active and regenerating cacao and pasture plots) and high values (old-growth forest plots). c) The percentage of forest 100 m around the centroid of the plots is not significantly correlated with elevation (Pearson’s *r* = 0.12, *p* = 0.37), but the percentage of forest d) 500 m (Pearson’s *r* = 0.26, *p* = 0.04) and e) 1 km (Pearson’s *r* = 0.44, *p* = 3E-04) around the centroid of the plots are. The percentage of forest at f) 100 m (Pearson’s *r* = 0.81, *p* = 1.5E-15), g) 500 m (Pearson’s *r* = 0.57, *p* = 1.9E-06), and h) 1 km (Pearson’s *r* = 0.38, *p* = 0.002) around the centroid of the plots are significantly correlated with square-root transformed regeneration time. Old-growth forest (OG) plots were not included in the analyses (f, g, h) because their time without human intervention is unknown and were used only as a reference. Dark circles in all boxplots represent mean values.

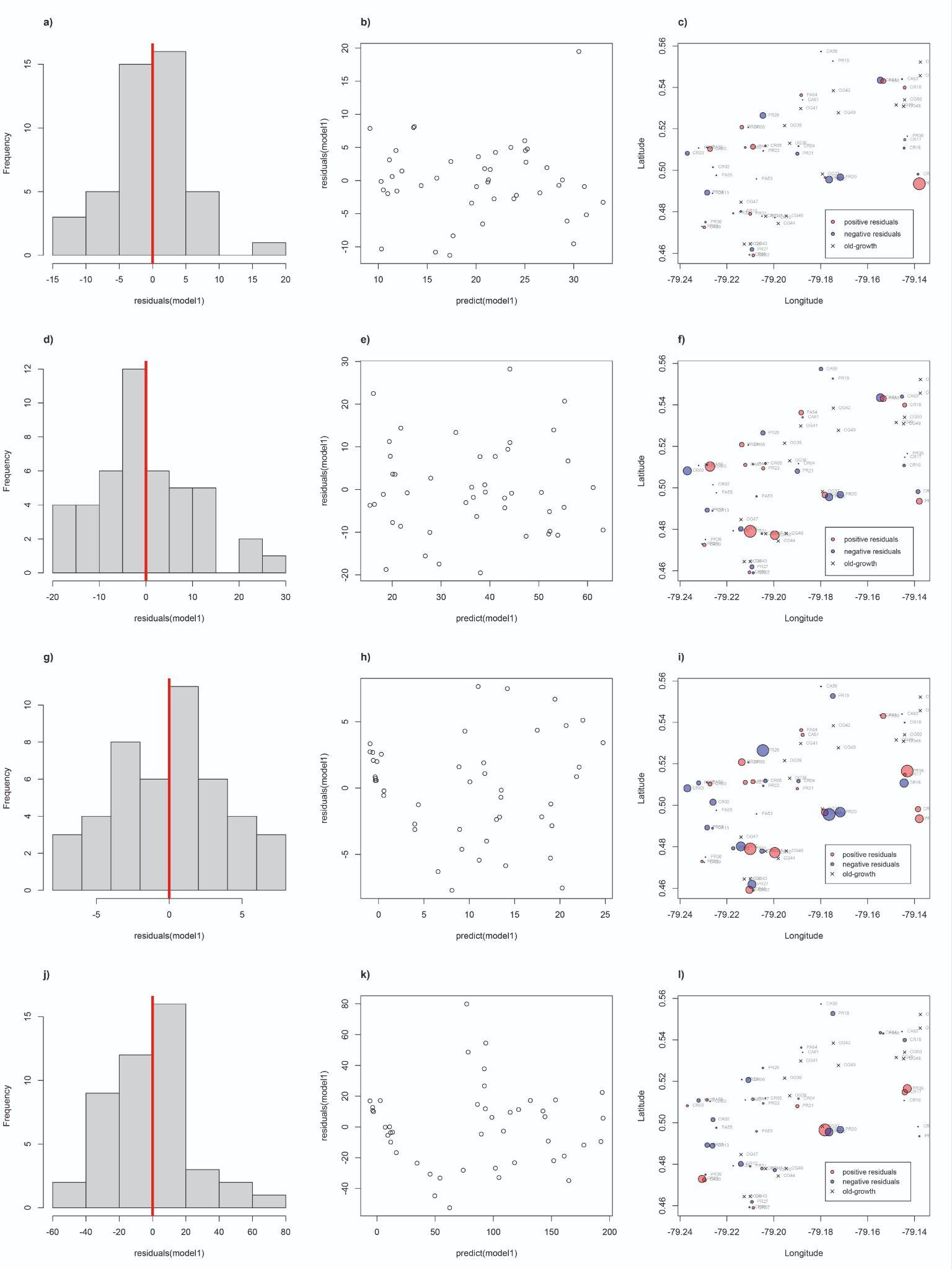

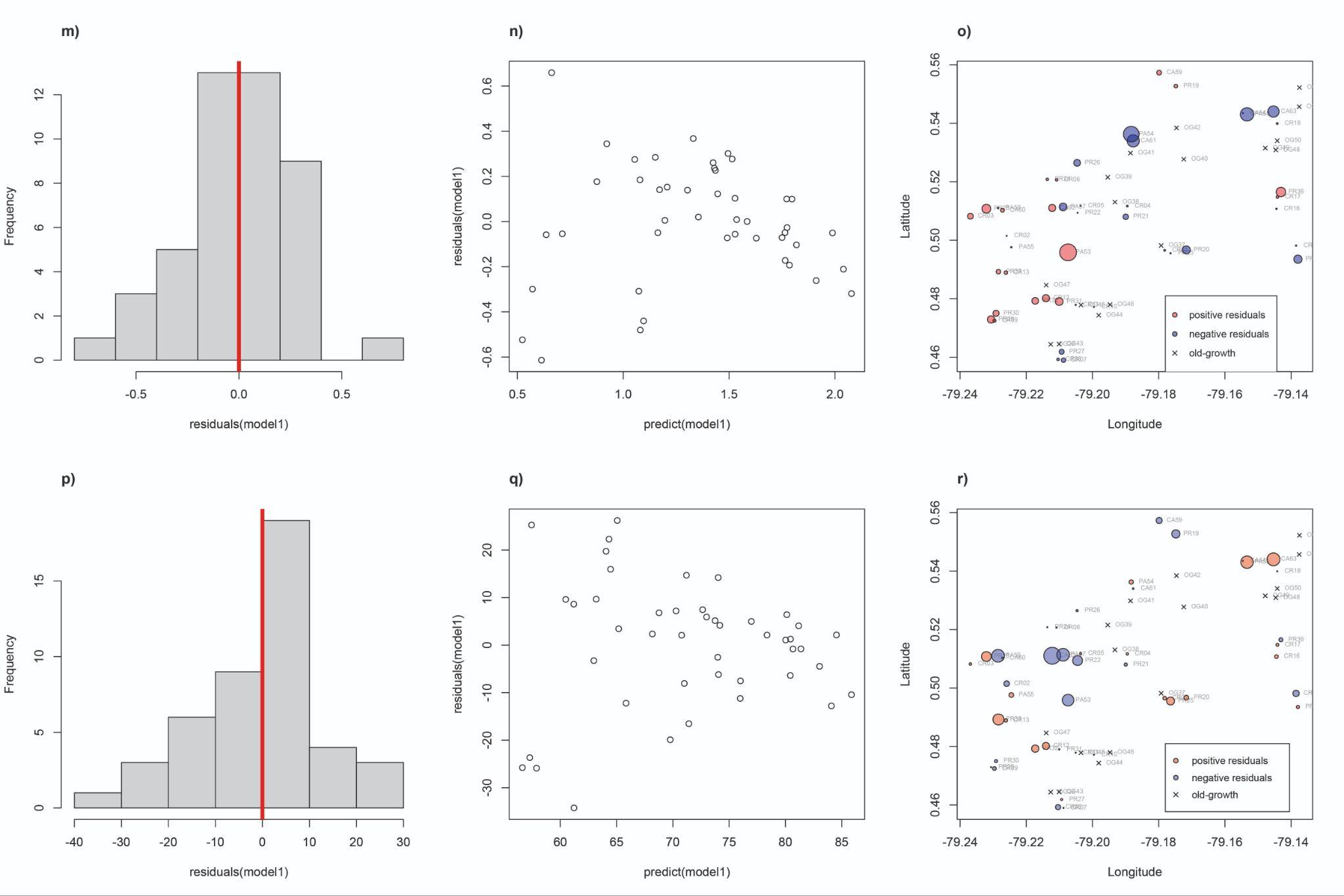

**Figure S3.** Linear mixed-effects model validation for forest structure variables. Histogram of the residuals, scatter plot of the fitted and residual values, and geographic distribution of the residuals of the models analyzed with maximum tree height (a, b, c), maximum tree DBH (d, e, f), basal area (g, h, i), number of stems(j, k, l), vertical vegetation heterogeneity (m, n, o), and reversed total site factor (TSF; p, q, r) as response variables. Histograms of the residuals (a, d, g, j, m, p) visually follow a normal distribution. The fitted and residual values of the models are not significantly correlated (b, e, h, k, n, q; all Pearson's *r* < 0.001, all *p* = 1). Spatial autocorrelation was only detected for maximum tree DBH (*p* < 0.05, f) and vertical vegetation heterogeneity (*p* < 0.05, o). Maximum tree height, maximum tree DBH, basal area, and number of stems were estimated for wild trees with DBH ≥ 7.95 cm. Reversed TSF was calculated as 1-TSF.

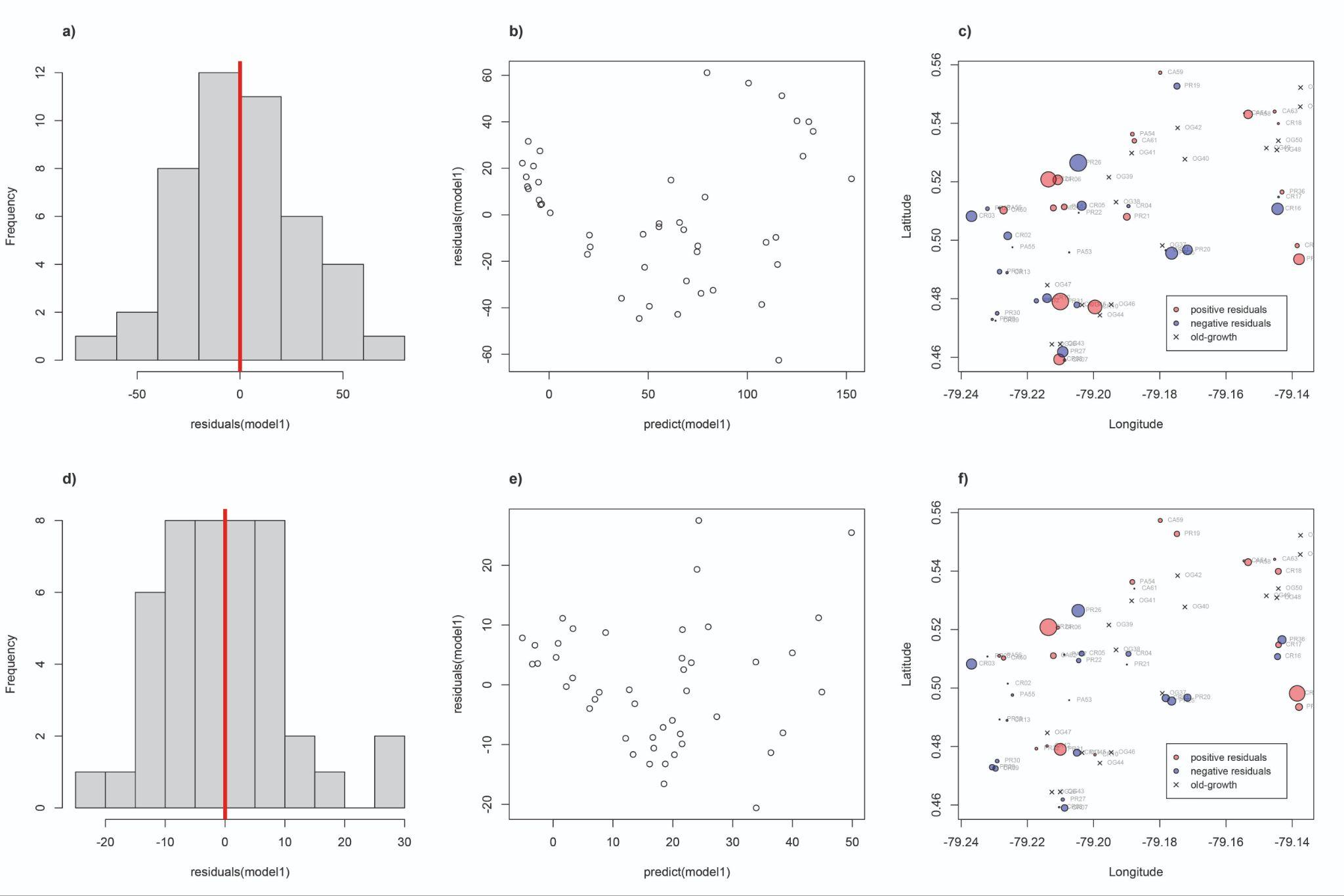

**Figure S4.** Linear mixed-effects model validation for aboveground biomass (AGB) and tree species richness. Histogram of the residuals, scatter plot of the fitted and residual values, and geographic distribution of the residuals of the models analyzed with AGB (a, b, c) and tree species richness (d, e, f) as response variables. Histograms of the residuals (a, d) visually follow a normal distribution. The fitted and residual values of the models are not correlated (b, e; Pearson's *r* < 0.001, *p* = 1). Spatial correlation was not observed (c, f; *p* > 0.05). AGB estimated using species-specific wood density along with DBH and height data, and rarefied tree species richness estimated with a coverage of 0.75 were estimated for wild trees with DBH ≥ 7.95 cm.

**Supplementary Tables**

**Table S1**. Baseline data on the 62 plots included in the study design.

| **Plot_ID** | **Treatment** | **Legacy** | **Reg_year** | **Duration_use** | **Latitude** | **Longitude** | **Elevation** |
| --- | --- | --- | --- | --- | --- | --- | --- |
| CA59 | cacao active | cacao | NA | 11 | 0.55731 | -79.1799 | 165 |
| CA60 | cacao active | cacao | NA | 23 | 0.510266 | -79.2272 | 440 |
| CA61 | cacao active | cacao | NA | 8 | 0.534001 | -79.1877 | 324 |
| CA62 | cacao active | cacao | NA | 6 | 0.511049 | -79.2122 | 372 |
| CA63 | cacao active | cacao | NA | 14 | 0.544018 | -79.1453 | 240 |
| CA64 | cacao active | cacao | NA | 23 | 0.543481 | -79.1546 | 256 |
| CR01 | cacao regeneration | cacao | 2017 | 17 | 0.496548 | -79.1782 | 560 |
| CR02 | cacao regeneration | cacao | 2015 | 15 | 0.501505 | -79.2259 | 400 |
| CR03 | cacao regeneration | cacao | 2018 | 6 | 0.508212 | -79.2369 | 274 |
| CR04 | cacao regeneration | cacao | 2002 | 7 | 0.511679 | -79.1895 | 560 |
| CR05 | cacao regeneration | cacao | 2011 | 13 | 0.511784 | -79.2036 | 377 |
| CR06 | cacao regeneration | cacao | 1990 | 3 | 0.520662 | -79.2108 | 397 |
| CR07 | cacao regeneration | cacao | 2000 | 30 | 0.458986 | -79.2088 | 168 |
| CR08 | cacao regeneration | cacao | 1996 | 26 | 0.459253 | -79.2104 | 172 |
| CR09 | cacao regeneration | cacao | 2017 | 6 | 0.472474 | -79.2296 | 159 |
| CR10 | cacao regeneration | cacao | 1999 | 9 | 0.477175 | -79.1996 | 347 |
| CR11 | cacao regeneration | cacao | 2000 | 5 | 0.477893 | -79.2051 | 372 |
| CR12 | cacao regeneration | cacao | 2008 | 7 | 0.480185 | -79.2141 | 324 |
| CR13 | cacao regeneration | cacao | 2022 | 7 | 0.488923 | -79.2262 | 366 |
| CR14 | cacao regeneration | cacao | 1985 | 5 | 0.49816 | -79.1385 | 529 |
| CR16 | cacao regeneration | cacao | 2015 | 15 | 0.510746 | -79.1444 | 479 |
| CR17 | cacao regeneration | cacao | 2014 | 14 | 0.51478 | -79.1441 | 450 |
| CR18 | cacao regeneration | cacao | 2013 | 10 | 0.539918 | -79.1442 | 331 |
| OG28 | old-growth | old-growth | NA | NA | 0.464406 | -79.2126 | 224 |
| OG37 | old-growth | old-growth | NA | NA | 0.498192 | -79.1792 | 550 |
| OG38 | old-growth | old-growth | NA | NA | 0.513057 | -79.1932 | 535 |
| OG39 | old-growth | old-growth | NA | NA | 0.521552 | -79.1954 | 576 |
| OG40 | old-growth | old-growth | NA | NA | 0.527723 | -79.1724 | 531 |
| OG41 | old-growth | old-growth | NA | NA | 0.529813 | -79.1885 | 389 |
| OG42 | old-growth | old-growth | NA | NA | 0.53843 | -79.1746 | 356 |
| OG43 | old-growth | old-growth | NA | NA | 0.464478 | -79.2101 | 238 |
| OG44 | old-growth | old-growth | NA | NA | 0.474349 | -79.1981 | 339 |
| OG45 | old-growth | old-growth | NA | NA | 0.477861 | -79.2035 | 362 |
| OG46 | old-growth | old-growth | NA | NA | 0.477978 | -79.1947 | 288 |
| OG47 | old-growth | old-growth | NA | NA | 0.484658 | -79.214 | 355 |
| OG48 | old-growth | old-growth | NA | NA | 0.53089 | -79.1446 | 299 |
| OG49 | old-growth | old-growth | NA | NA | 0.531535 | -79.1478 | 310 |
| OG50 | old-growth | old-growth | NA | NA | 0.53403 | -79.1442 | 316 |
| OG51 | old-growth | old-growth | NA | NA | 0.552232 | -79.1375 | 228 |
| OG52 | old-growth | old-growth | NA | NA | 0.545679 | -79.1376 | 225 |
| PA53 | pasture active | pasture | NA | 23 | 0.495852 | -79.2074 | 510 |
| PA54 | pasture active | pasture | NA | 23 | 0.536269 | -79.1883 | 207 |
| PA55 | pasture active | pasture | NA | 19 | 0.497611 | -79.2245 | 426 |
| PA56 | pasture active | pasture | NA | 23 | 0.511023 | -79.2285 | 430 |
| PA57 | pasture active | pasture | NA | 6 | 0.511392 | -79.2089 | 375 |
| PA58 | pasture active | pasture | NA | 7 | 0.543056 | -79.1533 | 243 |
| PR19 | pasture regeneration | pasture | 2017 | 17 | 0.552719 | -79.1748 | 177 |
| PR20 | pasture regeneration | pasture | 2019 | 19 | 0.496703 | -79.1717 | 540 |
| PR21 | pasture regeneration | pasture | 1987 | 1 | 0.508024 | -79.19 | 560 |
| PR22 | pasture regeneration | pasture | 2011 | 13 | 0.509393 | -79.2045 | 400 |
| PR23 | pasture regeneration | pasture | 2022 | 22 | 0.51078 | -79.2321 | 415 |
| PR24 | pasture regeneration | pasture | 2002 | 22 | 0.52082 | -79.2137 | 381 |
| PR25 | pasture regeneration | pasture | 2017 | 17 | 0.495558 | -79.1764 | 544 |
| PR26 | pasture regeneration | pasture | 1995 | 8 | 0.526466 | -79.2046 | 325 |
| PR27 | pasture regeneration | pasture | 1996 | 7 | 0.461851 | -79.2093 | 214 |
| PR29 | pasture regeneration | pasture | 2017 | 10 | 0.47292 | -79.2306 | 173 |
| PR30 | pasture regeneration | pasture | 2008 | 6 | 0.475008 | -79.2291 | 219 |
| PR31 | pasture regeneration | pasture | 2007 | 6 | 0.479026 | -79.2101 | 316 |
| PR32 | pasture regeneration | pasture | 2007 | 6 | 0.479275 | -79.2173 | 339 |
| PR33 | pasture regeneration | pasture | 2022 | 7 | 0.489245 | -79.2284 | 311 |
| PR34 | pasture regeneration | pasture | 1985 | 5 | 0.493509 | -79.1379 | 615 |
| PR36 | pasture regeneration | pasture | 2014 | 14 | 0.516501 | -79.1431 | 380 |

Continuation…

| **Plot_ID** | **Patch_ha** | **Distance** | **Forest_100m** | **Forest_500m** | **Forest_1km** | **Reversed_TSF** | **VVH** |
| --- | --- | --- | --- | --- | --- | --- | --- |
| CA59 | 18.47498 | 66.69162 | 4.263157 | 24.88893 | 33.47445 | 53.61905 | 1.263654 |
| CA60 | 2.371524 | 17.96248 | 13.3244 | 59.39237 | 71.93695 | 68.61905 | 1.359811 |
| CA61 | 3.916493 | 54.99201 | 7.231092 | 57.51965 | 79.7493 | 59.71429 | 0.600166 |
| CA62 | 14.87674 | 131.6154 | 0 | 23.52259 | 27.52259 | 26.96429 | 1.329297 |
| CA63 | 0.843782 | 143.8804 | 0 | 56.39467 | 83.18785 | 91.28571 | 0.655993 |
| CA64 | 5.376492 | 35.07122 | 16.06207 | 58.25824 | 67.55275 | 70.52381 | 1.201957 |
| CR01 | 0.492453 | 31.84109 | 23.43878 | 53.59181 | 76.82499 | 77.47619 | 1.418911 |
| CR02 | 0.946003 | 36.42346 | 56.05529 | 89.32652 | 89.5991 | 64.7619 | 1.585347 |
| CR03 | 1.458779 | 33.08483 | 53.06817 | 76.82607 | 73.73364 | 78.90476 | 1.661873 |
| CR04 | 1.861999 | 29.41603 | 54.65495 | 96.27259 | 97.12514 | 81.95238 | 1.678686 |
| CR05 | 4.790451 | 65.63923 | 10.17473 | 13.65745 | 41.15905 | 78.33333 | 1.544394 |
| CR06 | 0.643334 | 28.96937 | 34.34829 | 36.61514 | 41.12593 | 81.71429 | 1.896237 |
| CR07 | 3.75544 | 114.9121 | 0 | 57.12641 | 55.37701 | 86.66667 | 1.59261 |
| CR08 | 5.70617 | 49.9914 | 13.81523 | 65.87007 | 53.20305 | 75.42857 | 1.71472 |
| CR09 | 1.06906 | 39.94556 | 51.39917 | 43.9749 | 32.42567 | 68.47619 | 1.568565 |
| CR10 | 1.550152 | 25.04409 | 51.5854 | 91.85387 | 96.64203 | 80.57143 | 1.746987 |
| CR11 | 1.55386 | 36.46924 | 63.10319 | 91.32184 | 89.96987 | 79.85714 | 1.715221 |
| CR12 | 4.283623 | 157.6304 | 0 | 19.51279 | 74.76795 | 88.2381 | 1.79141 |
| CR13 | 3.211388 | 38.53007 | 52.54433 | 82.12563 | 77.95376 | 75.57143 | 1.442341 |
| CR14 | 0.526505 | 22.71859 | 83.21892 | 96.28904 | 98.61818 | 71.28571 | 1.937764 |
| CR16 | 0.818761 | 43.87424 | 29.00042 | 65.91047 | 84.5492 | 80.09524 | 1.472184 |
| CR17 | 1.297197 | 49.1178 | 24.23908 | 59.92649 | 86.1844 | 78.90476 | 1.630824 |
| CR18 | 0.275992 | 25.59526 | 74.63726 | 66.66145 | 85.03603 | 80.47619 | 1.554713 |
| OG28 | NA | 85.20277 | 98.82799 | 53.9589 | 64.95801 | 84 | 1.757592 |
| OG37 | NA | 122.8746 | 99.99999 | 66.80653 | 75.01316 | 79.04762 | 1.924961 |
| OG38 | NA | 403.3539 | 99.99998 | 96.97967 | 87.41406 | 77.09524 | 1.784249 |
| OG39 | NA | 780.6852 | 99.99998 | 100 | 96.66099 | 88.19048 | 1.883757 |
| OG40 | NA | 723.9293 | 99.99999 | 100 | 98.85801 | 92.71429 | 1.814715 |
| OG41 | NA | 214.0048 | 99.99999 | 94.62245 | 86.31303 | 82.42857 | 1.931368 |
| OG42 | NA | 188.4645 | 99.99998 | 83.70179 | 76.33876 | 80.71429 | 1.799954 |
| OG43 | NA | 147.8222 | 99.99998 | 78.76785 | 68.6117 | 86.09524 | 1.889783 |
| OG44 | NA | 302.6796 | 99.99996 | 97.73519 | 93.48595 | 70.38095 | 1.943601 |
| OG45 | NA | 35.81112 | 72.50359 | 89.50271 | 93.17093 | 90.52381 | 1.855436 |
| OG46 | NA | 463.8969 | 99.99998 | 99.65079 | 98.09357 | 81.14286 | 1.850355 |
| OG47 | NA | 82.05557 | 95.58186 | 65.33918 | 74.88779 | 82.38095 | 1.933311 |
| OG48 | NA | 942.6327 | 99.99998 | 100 | 99.82211 | 82.47619 | 1.830057 |
| OG49 | NA | 858.3927 | 99.99997 | 99.99999 | 98.937 | 87.71429 | 1.897939 |
| OG50 | NA | 607.9662 | 99.99999 | 100 | 93.20945 | 77.66667 | 1.834539 |
| OG51 | NA | 413.9922 | 99.99997 | 96.9384 | 82.84373 | 81.28571 | 1.869189 |
| OG52 | NA | 221.0001 | 99.99998 | 94.60685 | 87.53399 | 75.14286 | 1.892563 |
| PA53 | 0.102569 | 71.47299 | 8.973976 | 81.89321 | 87.27756 | 33.61905 | 1.321049 |
| PA54 | 10.67252 | 92.24954 | 0.574225 | 42.57853 | 77.3821 | 69.82143 | 0 |
| PA55 | 4.69022 | 8.485378 | 44.91552 | 89.31478 | 91.95721 | 70.09524 | 0.657925 |
| PA56 | 3.038455 | 85.73178 | 1.31309 | 53.80376 | 74.24558 | 32 | 0.577849 |
| PA57 | 11.95958 | 16.37455 | 24.61295 | 21.70184 | 24.4259 | 30.85714 | 0.272248 |
| PA58 | 24.17893 | 40.25524 | 6.850847 | 60.46259 | 69.02453 | 82.71429 | 0 |
| PR19 | 6.554911 | 56.7706 | 17.85128 | 33.78026 | 37.13616 | 54.89286 | 1.31255 |
| PR20 | 22.17156 | 142.7085 | 0 | 46.54237 | 73.48573 | 72.85714 | 0.765502 |
| PR21 | 1.064727 | 21.16522 | 72.21343 | 96.27259 | 95.66122 | 74.04762 | 1.829033 |
| PR22 | 7.753099 | 105.8189 | 0 | 36.82177 | 42.30408 | 49.85714 | 1.375103 |
| PR23 | 1.687798 | 30.92214 | 28.85542 | 62.05453 | 76.83124 | 83.80952 | 1.266398 |
| PR24 | 5.074808 | 22.19631 | 43.01681 | 37.42918 | 38.76569 | 81.09524 | 1.871864 |
| PR25 | 25.51887 | 201.5191 | 0 | 36.22153 | 78.227 | 80.42857 | 1.113993 |
| PR26 | 0.780134 | 25.02999 | 75.5212 | 90.06834 | 84.31897 | 78.57143 | 1.649689 |
| PR27 | 5.227 | 50.19792 | 38.68863 | 67.25883 | 64.5287 | 85.2381 | 1.59205 |
| PR29 | 1.237951 | 36.91717 | 29.46513 | 33.58899 | 29.96284 | 72.85714 | 1.435762 |
| PR30 | 5.068008 | 41.53222 | 19.40616 | 49.17466 | 41.9289 | 67.85714 | 1.666489 |
| PR31 | 27.54875 | 39.3488 | 30.6229 | 66.68842 | 75.4393 | 71.42857 | 1.796216 |
| PR32 | 46.70698 | 174.4493 | 0 | 33.64306 | 69.16992 | 85.90476 | 1.685077 |
| PR33 | 1.212158 | 23.11586 | 56.1268 | 73.53113 | 69.99333 | 86.61905 | 1.052204 |
| PR34 | 3.267871 | 24.54014 | 58.78685 | 95.75187 | 88.80019 | 86.52381 | 1.758754 |
| PR36 | 20.53949 | 35.06538 | 32.43683 | 66.12812 | 86.5253 | 62.95238 | 1.698149 |

**Table S2**. Summary of the *Reassembly* plots (*N* = 62) by land-use type and regeneration age across a 200 km^2^ area in northwestern Ecuador. Patch size was calculated for distinct areas of continuous land use around each 50×50m plot. Distance to the nearest forest edge was calculated from the plot centroid excluding all known forest disturbances <40 yrs (some areas include older selective logging or other disturbances). Forest cover was calculated as the percentage of the total forest area within different radii buffers around the plot centroid. Regeneration areas ≤15 yrs (Regen I) were visible on optical imagery whereas local knowledge, ground-truthing, and historic Landsat were used to classify regeneration >15 yrs (Regen II). SE = standard error.

| **Land use** |  | **Agriculture** | | **Regen I, 0-15 yrs** | | **Regen II, 16-38 yrs** | | **Old-growth** |
| --- | --- | --- | --- | --- | --- | --- | --- | --- |
|  |  | **Pasture** | **Cacao** | **Pasture** | **Cacao** | **Pasture** | **Cacao** | **Forest** |
| **(number of plots)** |  | **(6)** | **(6)** | **(8)** | **(9)** | **(8)** | **(8)** | **(17)** |
| Elevation (m) | Mean | 365 | 300 | 368 | 377 | 371 | 359 | 360 |
|  | SE | 48 | 40 | 50 | 39 | 52 | 51 | 29 |
|  | Min | 207 | 165 | 173 | 159 | 214 | 168 | 224 |
|  | Max | 510 | 440 | 544 | 560 | 615 | 560 | 576 |
| Patch size (ha) | Mean | 9.1 | 7.6 | 10.8 | 1.6 | 11.8 | 2.5 | - |
|  | SE | 3.5 | 3 | 3.6 | 0.5 | 5.8 | 0.7 | - |
|  | Min | 0.1 | 0.8 | 1.2 | 0.3 | 0.8 | 0.5 | - |
|  | Max | 24.2 | 18.5 | 25.5 | 4.8 | 46.7 | 5.7 | - |
| Distance (m) | Mean | 52 | 75 | 79 | 40 | 50 | 58 | 388 |
| forest edge | SE | 15 | 21 | 23 | 4 | 18 | 18 | 72 |
|  | Min | 8 | 18 | 23 | 26 | 21 | 23 | 36 |
|  | Max | 92 | 144 | 202 | 66 | 174 | 158 | 943 |
| **Forest cover (%)** | | | | | | | | |
| 100 m radius | Mean | 15 | 7 | 21 | 42 | 42 | 38 | 98 |
|  | SE | 7 | 3 | 7 | 7 | 9 | 11 | 2 |
|  | Min | 1 | 0 | 0 | 10 | 0 | 0 | 73 |
|  | Max | 45 | 16 | 56 | 75 | 76 | 83 | 100 |
| 500 m radius | Mean | 58 | 47 | 49 | 61 | 67 | 69 | 89 |
|  | SE | 10 | 7 | 6 | 8 | 9 | 10 | 4 |
|  | Min | 22 | 24 | 34 | 14 | 34 | 20 | 54 |
|  | Max | 89 | 59 | 74 | 89 | 96 | 96 | 100 |
| 1 km radius | Mean | 71 | 61 | 62 | 72 | 70 | 76 | 87 |
|  | SE | 10 | 10 | 8 | 7 | 7 | 8 | 3 |
|  | Min | 24 | 28 | 30 | 32 | 39 | 41 | 65 |
|  | Max | 92 | 83 | 87 | 90 | 96 | 99 | 100 |

**Table S3.** Pearson correlation coefficients among elevation and forest cover metrics for Reassembly plots (*N* = 62) located across a 200 km^2^ area in northwestern Ecuador. Forest cover within 100 m was highly correlated with distance to the forest edge but minimally correlated with forest cover within 1 km.

| **Metric** | **Age^1^** | **Elevation** | **Patch size** | **Distance forest** | **Distance edge** | **Forest 100m** | **Forest 500m** |
| --- | --- | --- | --- | --- | --- | --- | --- |
| Elevation | -0.18 | - | - | - | - | - | - |
| Patch size | 0.14 | -0.05 | - | - | - | - | - |
| Distance forest | 0.21 | -0.03 | 0.61 | - | - | - | - |
| Distance edge | - | 0.07 | -0.27 | -0.42 | - | - | - |
| Forest 100 m | -0.35 | 0.12 | -0.53 | -0.78 | 0.64 | - | - |
| Forest 500 m | -0.39 | 0.26 | -0.47 | -0.63 | 0.54 | 0.76 | - |
| Forest 1 km | -0.2 | 0.44 | -0.26 | -0.29 | 0.43 | 0.51 | 0.82 |

^1^Correlation with regeneration age for chronosequence plots (*N* = 33).

**Table S4.** Results on forest structure, aboveground biomass (AGB), and species richness in the 62 plots included in the study design. AGB estimates for the three datasets were highly correlated between the two methods used (all Pearson's *r* > 0.99, all *p* < 0.001, all n = 62 plots). Tree species richness for the three datasets was also highly correlated between the two methods used (all Pearson's *r* = 0.89–0.92, all *p* < 0.001, all n = 62 plots).

| **Plot_ID** | **Max_tree_height** | **Max_tree_DBH** | **Number_stems** | **AGB_all** | **AGB_wild** | **AGB_dbh10** |
| --- | --- | --- | --- | --- | --- | --- |
| CA59 | 10.32 | 13.10801 | 8 | 2.787576 | 1.135715 | 0.921353 |
| CA60 | 17.02 | 38.64285 | 10 | 22.89548 | 22.81815 | 22.68901 |
| CA61 | 12.02 | 23.72047 | 11 | 12.65336 | 4.732821 | 4.704484 |
| CA62 | 14.26 | 27.40014 | 7 | 13.07772 | 8.706744 | 8.706744 |
| CA63 | 9 | 12.34248 | 4 | 11.46519 | 0.70952 | 0.625191 |
| CA64 | 0 | 0 | 0 | 0.170411 | 0 | 0 |
| CR01 | 20.3 | 46.40962 | 157 | 50.78292 | 50.25193 | 45.92816 |
| CR02 | 22.68 | 38.4646 | 75 | 48.25308 | 40.79876 | 40.39214 |
| CR03 | 14.12 | 18.53838 | 101 | 11.19531 | 11.19531 | 9.39489 |
| CR04 | 24.68 | 49.87284 | 153 | 98.10544 | 97.60983 | 93.70121 |
| CR05 | 19.08 | 38.77654 | 105 | 47.00866 | 42.84161 | 41.00429 |
| CR06 | 30.18 | 61.62485 | 130 | 174.2045 | 169.1034 | 167.1477 |
| CR07 | 29.58 | 47.03987 | 149 | 108.8982 | 104.6958 | 101.7087 |
| CR08 | 29.2 | 62.71347 | 138 | 167.0856 | 165.424 | 162.5021 |
| CR09 | 26.24 | 48.85424 | 119 | 58.5746 | 51.72222 | 48.47263 |
| CR10 | 27.76 | 76.0061 | 130 | 170.7984 | 168.6575 | 165.208 |
| CR11 | 28.94 | 51.01239 | 171 | 94.23184 | 93.85532 | 89.9804 |
| CR12 | 21.12 | 36.47198 | 72 | 51.95116 | 50.23783 | 47.78175 |
| CR13 | 16.36 | 30.55141 | 21 | 17.02834 | 11.58909 | 11.38873 |
| CR14 | 29.72 | 53.69892 | 200 | 168.0343 | 168.0343 | 162.7992 |
| CR16 | 16.14 | 30.99704 | 85 | 23.64869 | 22.0127 | 20.066 |
| CR17 | 23.84 | 40.06888 | 130 | 62.35505 | 61.51017 | 59.34792 |
| CR18 | 28.56 | 53.10687 | 95 | 88.06754 | 86.26249 | 84.16829 |
| OG28 | 37.26 | 100.8343 | 236 | 411.4048 | 411.4048 | 404.6945 |
| OG37 | 33.96 | 105.7936 | 195 | 482.5621 | 482.5621 | 477.2435 |
| OG38 | 27.56 | 62.57978 | 148 | 167.7737 | 167.7737 | 164.0086 |
| OG39 | 32.98 | 71.63888 | 169 | 331.322 | 331.322 | 328.3539 |
| OG40 | 38.68 | 114.6871 | 199 | 568.3411 | 568.3411 | 563.9639 |
| OG41 | 39.74 | 118.0803 | 149 | 554.0874 | 554.0874 | 550.5309 |
| OG42 | 42.3 | 89.04408 | 203 | 269.0316 | 268.9615 | 265.063 |
| OG43 | 36.62 | 129.826 | 231 | 506.1702 | 506.1702 | 495.1548 |
| OG44 | 34.78 | 99.44646 | 198 | 387.6717 | 387.2874 | 381.961 |
| OG45 | 31.28 | 86.79681 | 187 | 325.7297 | 325.7297 | 322.3715 |
| OG46 | 33.96 | 86.30025 | 155 | 300.6303 | 300.6303 | 297.9593 |
| OG47 | 33.74 | 58.88738 | 158 | 171.0831 | 171.0831 | 167.9662 |
| OG48 | 39.48 | 106.6976 | 161 | 420.4536 | 420.4536 | 416.9278 |
| OG49 | 37.04 | 91.0367 | 139 | 324.1274 | 324.1274 | 322.9785 |
| OG50 | 30.2 | 74.93021 | 200 | 191.4275 | 191.4275 | 188.3268 |
| OG51 | 38.24 | 117.2336 | 161 | 429.8627 | 429.8627 | 426.8597 |
| OG52 | 36.4 | 84.03388 | 143 | 254.1117 | 254.1117 | 251.4532 |
| PA53 | 10.16 | 11.75201 | 9 | 0.841214 | 0.841214 | 0.484613 |
| PA54 | 16.34 | 30.65327 | 7 | 9.209642 | 8.693403 | 8.693403 |
| PA55 | 13.85 | 17.03755 | 2 | 1.708514 | 1.364073 | 1.364073 |
| PA56 | 9.1 | 12.87246 | 11 | 1.238515 | 1.238515 | 1.114699 |
| PA57 | 21.6 | 24.10881 | 20 | 13.04157 | 13.04157 | 13.04157 |
| PA58 | 21.84 | 36.16003 | 7 | 21.04557 | 21.04557 | 21.04557 |
| PR19 | 19.14 | 32.03473 | 46 | 25.5894 | 25.5894 | 24.32751 |
| PR20 | 5.04 | 11.21725 | 5 | 0.572228 | 0.572228 | 0.519884 |
| PR21 | 26.12 | 43.22015 | 216 | 153.3002 | 153.3002 | 148.7729 |
| PR22 | 23.12 | 45.69661 | 105 | 62.45467 | 62.45467 | 59.05338 |
| PR23 | 13.62 | 22.29444 | 11 | 6.892375 | 6.892375 | 6.609366 |
| PR24 | 31 | 55.13132 | 151 | 157.2817 | 157.2817 | 153.5645 |
| PR25 | 6.02 | 12.14672 | 10 | 0.854491 | 0.854491 | 0.646478 |
| PR26 | 20.44 | 41.63497 | 166 | 53.49908 | 53.31576 | 48.37087 |
| PR27 | 23.2 | 42.50077 | 142 | 68.64232 | 68.64232 | 64.42916 |
| PR29 | 21.38 | 34.78493 | 127 | 39.33975 | 38.97654 | 37.07925 |
| PR30 | 27.88 | 43.57029 | 124 | 61.60926 | 61.60926 | 56.38492 |
| PR31 | 30.02 | 72.34553 | 133 | 140.8203 | 140.8203 | 137.0599 |
| PR32 | 21.9 | 41.02381 | 106 | 58.81005 | 58.69513 | 55.2664 |
| PR33 | 9.3 | 17.66621 | 15 | 2.287332 | 2.287332 | 2.165729 |
| PR34 | 50 | 67.04885 | 183 | 171.0477 | 171.0477 | 167.0761 |
| PR36 | 20.98 | 36.80939 | 148 | 76.44878 | 76.40301 | 75.05918 |

Continuation…

| **Plot_ID** | **AGC_all** | **AGC_wild** | **AGC_dbh10** | **AGB_Chave**  **_all** | **AGB_Chave**  **_wild** | **AGB_Chave**  **_dbh10** | **basal_area**  **_wild** |
| --- | --- | --- | --- | --- | --- | --- | --- |
| CA59 | 1.315763 | 0.536059 | 0.434699 | 2.963818 | 1.245959 | 0.977181 | 0.36946 |
| CA60 | 10.83229 | 10.746 | 10.63063 | 17.49436 | 17.4489 | 17.31702 | 2.852707 |
| CA61 | 5.973175 | 2.283516 | 2.239463 | 11.1116 | 5.149392 | 5.073066 | 1.497741 |
| CA62 | 6.15256 | 4.125851 | 4.071205 | 11.79913 | 7.831022 | 7.831022 | 1.784474 |
| CA63 | 5.40575 | 0.338241 | 0.293361 | 10.85055 | 0.61869 | 0.556192 | 0.199608 |
| CA64 | 0.080797 | 0 | 0 | 0.104636 | 0 | 0 | 0 |
| CR01 | 23.9585 | 23.75938 | 21.75644 | 58.52803 | 58.3175 | 55.10106 | 13.80237 |
| CR02 | 22.76081 | 19.21374 | 19.04778 | 49.7765 | 41.50684 | 41.08213 | 7.909014 |
| CR03 | 5.28724 | 5.284054 | 4.412674 | 14.45157 | 14.45157 | 12.21236 | 4.560115 |
| CR04 | 46.43287 | 45.89667 | 43.95894 | 88.1489 | 87.64502 | 84.73392 | 15.81634 |
| CR05 | 22.20553 | 20.20158 | 19.38226 | 46.78443 | 42.30087 | 41.00205 | 10.63165 |
| CR06 | 81.92914 | 79.3329 | 78.64622 | 168.489 | 162.3497 | 160.6709 | 22.68686 |
| CR07 | 51.41077 | 49.11734 | 47.67923 | 105.4858 | 101.6023 | 99.34045 | 17.79182 |
| CR08 | 78.52777 | 77.49689 | 76.50774 | 175.6228 | 173.8243 | 171.6413 | 25.39069 |
| CR09 | 27.56225 | 24.32972 | 22.85562 | 64.1061 | 56.04594 | 52.98632 | 10.53146 |
| CR10 | 80.44356 | 79.50572 | 78.08936 | 177.7916 | 175.6487 | 172.8045 | 26.14175 |
| CR11 | 44.47004 | 44.47425 | 42.30212 | 83.96874 | 83.59454 | 80.89874 | 16.27124 |
| CR12 | 24.39907 | 23.54787 | 22.48008 | 45.66284 | 44.29292 | 42.24527 | 8.156208 |
| CR13 | 8.020828 | 5.472408 | 5.336425 | 19.36646 | 13.77221 | 13.52242 | 3.12907 |
| CR14 | 79.37552 | 78.88502 | 76.61357 | 186.1096 | 186.1096 | 182.2965 | 28.17106 |
| CR16 | 11.05829 | 10.37793 | 9.439867 | 24.6314 | 22.78835 | 21.26706 | 5.651857 |
| CR17 | 29.39889 | 29.06329 | 27.96397 | 71.65873 | 70.79263 | 69.00845 | 13.49378 |
| CR18 | 41.46944 | 40.70924 | 39.50303 | 90.93297 | 89.07139 | 86.81003 | 12.78768 |
| OG28 | 194.0413 | 193.6601 | 190.3995 | 395.1632 | 395.1632 | 390.5251 | 46.18308 |
| OG37 | 226.4337 | 227.6329 | 224.6607 | 417.5831 | 417.5831 | 414.8614 | 48.18187 |
| OG38 | 78.70472 | 79.02666 | 77.36273 | 148.5405 | 148.5405 | 145.6062 | 22.37227 |
| OG39 | 155.4673 | 155.873 | 154.7234 | 277.9775 | 277.9775 | 275.8207 | 38.07224 |
| OG40 | 266.44 | 268.5961 | 265.3372 | 462.4956 | 462.4956 | 459.4253 | 49.26581 |
| OG41 | 261.0799 | 259.7253 | 259.1806 | 520.8736 | 520.8736 | 518.4464 | 51.21741 |
| OG42 | 127.9153 | 127.9847 | 125.0804 | 318.1286 | 318.0127 | 315.293 | 35.21305 |
| OG43 | 237.9154 | 238.7413 | 232.1851 | 476.3941 | 476.3941 | 470.5291 | 64.87576 |
| OG44 | 182.2385 | 182.5944 | 180.2508 | 326.1195 | 325.6272 | 321.4548 | 39.46105 |
| OG45 | 153.2857 | 153.4106 | 152.4527 | 299.6087 | 299.6087 | 295.9249 | 38.9399 |
| OG46 | 140.8239 | 141.5046 | 140.7605 | 266.746 | 266.746 | 263.2668 | 35.63245 |
| OG47 | 80.3299 | 80.47321 | 79.22419 | 163.5813 | 163.5813 | 161.252 | 23.4592 |
| OG48 | 197.5947 | 198.0519 | 195.389 | 394.6178 | 394.6178 | 392.206 | 41.83173 |
| OG49 | 153.1305 | 152.5035 | 153.0153 | 302.8575 | 302.8575 | 300.8325 | 34.70803 |
| OG50 | 90.00897 | 90.28279 | 88.73964 | 181.8225 | 181.8225 | 179.2477 | 28.94262 |
| OG51 | 200.6925 | 202.1184 | 200.5962 | 430.0826 | 430.0826 | 428.0443 | 43.07329 |
| OG52 | 120.8431 | 120.1975 | 118.874 | 271.8805 | 271.8805 | 269.3163 | 30.75148 |
| PA53 | 0.399213 | 0.400855 | 0.231165 | 1.006774 | 1.006774 | 0.632027 | 0.322622 |
| PA54 | 4.36886 | 4.101714 | 4.097203 | 8.688069 | 8.379248 | 8.379248 | 1.819972 |
| PA55 | 0.807417 | 0.633068 | 0.649766 | 2.043675 | 1.811445 | 1.811445 | 0.381475 |
| PA56 | 0.588595 | 0.582927 | 0.535595 | 1.204982 | 1.204982 | 1.116293 | 0.461552 |
| PA57 | 6.16793 | 6.144354 | 6.139463 | 12.1985 | 12.1985 | 12.1985 | 2.039714 |
| PA58 | 9.842398 | 9.838893 | 9.92132 | 17.00445 | 17.00445 | 17.00445 | 2.39961 |
| PR19 | 12.0048 | 12.10475 | 11.54125 | 24.22142 | 24.22142 | 23.37773 | 5.824567 |
| PR20 | 0.267909 | 0.268977 | 0.245108 | 0.403004 | 0.403004 | 0.36716 | 0.257943 |
| PR21 | 72.24867 | 72.10365 | 69.85782 | 139.5581 | 139.5581 | 135.9085 | 23.69835 |
| PR22 | 29.36151 | 29.47238 | 27.68943 | 57.31392 | 57.31392 | 54.97984 | 12.8028 |
| PR23 | 3.205963 | 3.258669 | 3.070761 | 5.526684 | 5.526684 | 5.243831 | 1.267664 |
| PR24 | 73.92553 | 74.20591 | 72.23382 | 150.8137 | 150.8137 | 147.886 | 21.85526 |
| PR25 | 0.404223 | 0.402226 | 0.303905 | 0.665638 | 0.665638 | 0.496258 | 0.375677 |
| PR26 | 25.17734 | 25.06151 | 22.74848 | 54.97632 | 54.87473 | 50.81133 | 12.71192 |
| PR27 | 32.21992 | 32.31323 | 30.35828 | 71.01658 | 71.01658 | 67.59827 | 13.66458 |
| PR29 | 18.48268 | 18.37006 | 17.41904 | 52.15807 | 51.78728 | 49.76014 | 10.46434 |
| PR30 | 28.97398 | 29.0127 | 26.55995 | 77.96909 | 77.96909 | 73.90516 | 13.36456 |
| PR31 | 66.44245 | 66.38297 | 64.72396 | 165.8786 | 165.8786 | 162.9684 | 21.69132 |
| PR32 | 27.81605 | 27.60214 | 25.96366 | 56.30017 | 56.2241 | 53.77012 | 11.15422 |
| PR33 | 1.079005 | 1.082285 | 1.020387 | 2.335817 | 2.335817 | 2.160826 | 0.869318 |
| PR34 | 80.42325 | 81.05442 | 79.58071 | 202.3533 | 202.3533 | 199.693 | 27.63492 |
| PR36 | 35.95562 | 35.96455 | 35.25774 | 100.8272 | 100.7994 | 99.39063 | 18.63135 |

Continuation…

| **Plot_ID** | **sr_all** | **sr_wild** | **sr_dbh10** | **sr_rar_all** | **sr_rar_wild** | **sr_rar_dbh10** |
| --- | --- | --- | --- | --- | --- | --- |
| CA59 | 5 | 4 | 3 | 2.812571 | 3.584374 | 3.5 |
| CA60 | 7 | 6 | 5 | 10.84091 | 7.775 | 6.0625 |
| CA61 | 8 | 3 | 3 | 6.447645 | 1.904779 | 2.142859 |
| CA62 | 10 | 6 | 6 | 11.84239 | 12.71429 | 12.71429 |
| CA63 | 4 | 1 | 1 | 1 | 1 | 1 |
| CA64 | 1 | 0 | 0 | 1 | 0 | 0 |
| CR01 | 21 | 20 | 18 | 8.993121 | 8.575623 | 8.932323 |
| CR02 | 29 | 28 | 27 | 11.93446 | 21.29383 | 22.16277 |
| CR03 | 7 | 7 | 5 | 1.950072 | 1.950072 | 1.71499 |
| CR04 | 48 | 46 | 39 | 31.47946 | 30.35512 | 32.23276 |
| CR05 | 28 | 26 | 21 | 10.35986 | 13.03396 | 10.23903 |
| CR06 | 49 | 48 | 43 | 27.03717 | 45.30457 | 61.9387 |
| CR07 | 21 | 19 | 19 | 7.035123 | 6.227121 | 6.513983 |
| CR08 | 41 | 40 | 36 | 21.30296 | 24.35888 | 25.66961 |
| CR09 | 21 | 19 | 16 | 6.118525 | 7.864476 | 6.85718 |
| CR10 | 50 | 48 | 35 | 36.02956 | 37.67001 | 28.3489 |
| CR11 | 47 | 45 | 39 | 27.39996 | 25.0312 | 25.95144 |
| CR12 | 33 | 31 | 27 | 26.92348 | 26.79167 | 26.26574 |
| CR13 | 12 | 10 | 9 | 1.39742 | 10.46591 | 9.143519 |
| CR14 | 80 | 80 | 60 | 75.29818 | 75.29818 | 57.94353 |
| CR16 | 21 | 20 | 17 | 8.890454 | 11.66669 | 10.75811 |
| CR17 | 43 | 42 | 31 | 27.21019 | 30.83521 | 20.47517 |
| CR18 | 41 | 40 | 33 | 26.91145 | 35.61027 | 33.21507 |
| OG28 | 63 | 63 | 56 | 34.38046 | 34.38046 | 37.6577 |
| OG37 | 66 | 66 | 54 | 52.81824 | 52.81824 | 43.52966 |
| OG38 | 64 | 64 | 53 | 63.33168 | 63.33168 | 57.4442 |
| OG39 | 64 | 64 | 57 | 56.42704 | 56.42704 | 58.53714 |
| OG40 | 73 | 73 | 61 | 63.82126 | 63.82126 | 65.71711 |
| OG41 | 68 | 68 | 57 | 89.40059 | 89.40059 | 95.41379 |
| OG42 | 90 | 89 | 69 | 99.76244 | 97.30078 | 67.84392 |
| OG43 | 76 | 76 | 62 | 56.85521 | 56.85521 | 56.02481 |
| OG44 | 85 | 84 | 76 | 94.15538 | 91.72494 | 135.1128 |
| OG45 | 90 | 90 | 71 | 140.6638 | 140.6638 | 134.0243 |
| OG46 | 72 | 72 | 56 | 98.43498 | 98.43498 | 73.09375 |
| OG47 | 71 | 71 | 63 | 77.54761 | 77.54761 | 90.00419 |
| OG48 | 61 | 61 | 54 | 50.89135 | 50.89135 | 48.70402 |
| OG49 | 61 | 61 | 53 | 61.2465 | 61.2465 | 57.50862 |
| OG50 | 78 | 78 | 64 | 67.99595 | 67.99595 | 57.84781 |
| OG51 | 50 | 50 | 43 | 34.01844 | 34.01844 | 30.2487 |
| OG52 | 64 | 64 | 55 | 83.77564 | 83.77564 | 82.99091 |
| PA53 | 4 | 4 | 1 | 4.416667 | 4.416667 | 1 |
| PA54 | 5 | 3 | 3 | 7.083333 | 2.704313 | 2.704313 |
| PA55 | 3 | 2 | 2 | 3.05 | 2.125 | 2.125 |
| PA56 | 5 | 5 | 4 | 5.090909 | 5.090909 | 4.416667 |
| PA57 | 5 | 5 | 5 | 4.56704 | 4.56704 | 4.56704 |
| PA58 | 6 | 6 | 6 | 12.71429 | 12.71429 | 12.71429 |
| PR19 | 18 | 18 | 17 | 17.50717 | 17.50717 | 27.44444 |
| PR20 | 2 | 2 | 2 | 1.709765 | 1.709765 | 1.263953 |
| PR21 | 65 | 65 | 56 | 43.68175 | 43.68175 | 40.39779 |
| PR22 | 24 | 24 | 19 | 11.25214 | 11.25214 | 9.799863 |
| PR23 | 7 | 7 | 5 | 6.453549 | 6.453549 | 5.553571 |
| PR24 | 60 | 60 | 45 | 51.81764 | 51.81764 | 40.49612 |
| PR25 | 4 | 4 | 3 | 2.874531 | 2.874531 | 2.512502 |
| PR26 | 34 | 32 | 26 | 14.2727 | 13.3372 | 13.82651 |
| PR27 | 37 | 37 | 31 | 22.0145 | 22.0145 | 22.05496 |
| PR29 | 15 | 14 | 13 | 4.310412 | 3.168796 | 2.104218 |
| PR30 | 30 | 30 | 25 | 13.97417 | 13.97417 | 15.47623 |
| PR31 | 50 | 50 | 38 | 43.3435 | 43.3435 | 36.4065 |
| PR32 | 38 | 37 | 35 | 26.96799 | 25.97878 | 33.66063 |
| PR33 | 9 | 9 | 8 | 11.9 | 11.9 | 10.04167 |
| PR34 | 68 | 68 | 62 | 55.55973 | 55.55973 | 56.70593 |
| PR36 | 19 | 18 | 17 | 5.674342 | 5.513443 | 5.025963 |

**Table S5.** Results of linear models. a) Maximum tree height, b) maximum tree DBH, c) basal area, d) number of stems, e) vertical vegetation heterogeneity, and f) reversed total site factor (TSF).

| **Max tree height - ANOVA type II error** | | | | |
| --- | --- | --- | --- | --- |
| **Variable** | **Sum Sq** | **Df** | ***F* value** | ***p*** |
| Elevation (masl) | 22.31 | 1 | 0.588 | 0.4479 |
| Forest 100 m around | 17.75 | 1 | 0.4678 | 0.4982 |
| Land-use legacy | 2.12 | 1 | 0.056 | 0.8143 |
| sqrt(Regeneration time) | 1275.6 | 1 | 3.36E+01 | **1.08E-06** |
| Duration use | 23.67 | 1 | 0.6237 | 0.4346 |
| Land-use legacy (pasture) : sqrt(Regeneration time) | 9.69 | 1 | 0.2554 | 0.6162 |
| Residuals | 1442 | 38 |  |  |
| **Max tree DBH - ANOVA type II error** | | | | |
| Elevation (masl) | 121.5 | 1 | 0.9048 | 0.3475 |
| Forest 100 m around | 10.8 | 1 | 0.0801 | 0.7787 |
| Land-use legacy | 128 | 1 | 0.9533 | 0.335 |
| sqrt(Regeneration time) | 5688.1 | 1 | 42.3711 | **1.14E-07** |
| Duration use | 47.8 | 1 | 0.3563 | 0.5541 |
| Land-use legacy (pasture) : sqrt(Regeneration time) | 88.5 | 1 | 0.6592 | 0.4219 |
| Residuals | 5101.3 | 38 |  |  |
| **Basal area - ANOVA type II error** | | | | |
| Elevation (masl) | 0.64 | 1 | 0.0377 | 0.8472 |
| Forest 100 m around | 12.87 | 1 | 0.7536 | 0.3908 |
| Land-use legacy | 7.97 | 1 | 0.4666 | 0.4987 |
| sqrt(Regeneration time) | 1967 | 1 | 115.1375 | **4.65E-13** |
| Duration use | 5.31 | 1 | 0.3111 | 0.5803 |
| Land-use legacy (pasture) : sqrt(Regeneration time) | 1.45 | 1 | 0.0848 | 0.7725 |
| Residuals | 649.18 | 38 |  |  |
| **Number of stems - ANOVA type II error** | | | | |
| Elevation (masl) | 84 | 1 | 0.1061 | 0.7464 |
| Forest 100 m around | 2969 | 1 | 3.7584 | 0.06 |
| Land-use legacy | 191 | 1 | 0.2417 | 0.6258 |
| sqrt(Regeneration time) | 101586 | 1 | 128.6072 | **9.25E-14** |
| Duration use | 13 | 1 | 0.0167 | 0.8977 |
| Land-use legacy (pasture) : sqrt(Regeneration time) | 662 | 1 | 0.8387 | 0.3656 |
| Residuals | 30016 | 38 |  |  |
| **Vertical vegetation heterogeneity - ANOVA type II error** | | | | |
| Elevation (masl) | 0.0055 | 1 | 0.0714 | 0.790689 |
| Forest 100 m around | 0.0969 | 1 | 1.2499 | 0.270599 |
| Land-use legacy | 0.6654 | 1 | 8.5816 | **0.005715** |
| sqrt(Regeneration time) | 4.1354 | 1 | 53.3366 | **9.60E-09** |
| Duration use | 0.0773 | 1 | 0.997 | 0.324358 |
| Land-use legacy (pasture) : sqrt(Regeneration time) | 0.5429 | 1 | 7.0016 | **0.011781** |
| Residuals | 2.9463 | 38 |  |  |
| **Reversed TSF - ANOVA type II error** | | | | |
| Elevation (masl) | 146.1 | 1 | 0.6922 | 0.410619 |
| Forest 100 m around | 108.4 | 1 | 0.5138 | 0.47789 |
| Land-use legacy | 175.5 | 1 | 0.8317 | 0.367518 |
| sqrt(Regeneration time) | 1729.5 | 1 | 8.1957 | **0.006796** |
| Duration use | 90.4 | 1 | 0.4283 | 0.51675 |
| Land-use legacy (pasture) : sqrt(Regeneration time) | 38.9 | 1 | 0.1843 | 0.67014 |
| Residuals | 8019 | 38 |  |  |
